# Structures of the human adult muscle-type nicotinic receptor in resting and desensitised states

**DOI:** 10.1101/2024.12.13.624530

**Authors:** Anna Li, Ashley C. W. Pike, Richard Webster, Susan Maxwell, Wei-Wei Liu, Gamma Chi, Jacqueline Palace, David Beeson, David B. Sauer, Yin Yao Dong

**Affiliations:** Nuffield Department of Clinical Neurosciences, University of Oxford, Oxford, UK; Centre for Medicines Discovery, Nuffield Department of Medicine, University of Oxford, Oxford, UK; Neurology Department, John Radcliffe Hospital, Oxford OX3 9DU, UK

**Author notes:** Corresponding contact (X: @Anna_Liiiiiiii). Corresponding and lead contact (X: @CMS_CDG_Oxford).

**Keywords:** Nicotinic receptor, pLGIC, congenital myasthenia, ion channel, electrophysiology, neuromuscular junction, cryo-EM, acetylcholine receptor

## Abstract

Muscle-type nicotinic acetylcholine receptor (AChR) is the key signalling molecule in neuromuscular junctions. Here we present the structures of full-length human adult receptors in complex with Fab35 in α-bungarotoxin (αBuTx)-bound resting states, and in acetylcholine (ACh)-bound desensitised states. In addition to identifying the conformational changes during recovery from desensitisation, we also used electrophysiology to probe the effects of eight previously unstudied AChR genetic variants found in congenital myasthenic syndrome (CMS) patients, revealing they cause either slow- or fast-channel CMS characterised by prolonged or abbreviated ion channel bursts. The combined kinetic and structural data offer a better understanding of both AChR state transition and the pathogenic mechanisms of disease variants.

## Introduction

AChR is a pentameric ligand-gated ion channel (pLGIC) found clustered on the post-synaptic membrane of neuromuscular junctions. Consisting of four different subunits (α, β, δ, and γ or ε), it is the most complex pLGIC by subunit heterogeneity. At the neuromuscular junction, binding of acetylcholine (ACh) drives the transition from resting to open state, which results in depolarisation of the membrane and eventually muscle contraction. Subsequent dissociation of ACh causes the receptor to return to the resting state. Occasionally, high ACh concentration may prolong the open state of AChR and prompt its transition to a non-conductive desensitised state, and this mechanism safeguards the post-synaptic membrane from overstimulation. In the neuromuscular junctions of developing mammals, AChR undergoes subunit-switching where its foetal γ subunit is substituted by an ε subunit, resulting in an adult-type receptor (α_2_βδε) with shorter burst duration and higher single-channel conductance^1,2^.

Abnormalities in neuromuscular signalling leads to myasthenia (fatigable muscle weakness), which can affect all voluntary muscles in the body. When respiratory muscles are affected, patients may be at risk of myasthenic crises, and in rare cases even death. Myasthenia can be caused by autoimmune attack in the case of myasthenia gravis, or genetic defects in proteins of the neuromuscular junction in the case of congenital myasthenic syndrome.

Prevalence of CMS varies between 2-22 per 1,000,000 people worldwide^3–6^, and is most commonly driven by mutation in AChR for half of all cases^7,8^. Variants in AChR genes can lead to CMS by three major pathogenic mechanisms – reduced receptor surface expression (AChR deficiency syndrome), prolonged AChR activity (slow channel congenital myasthenic syndromes, SCCMS), or abbreviated AChR activity (fast channel congenital myasthenic syndromes, FCCMS)^9^. Strategies for treating FCCMS, SCCMS and AChR deficiency syndrome are distinct and sometimes contraindicated, therefore a correct diagnosis is vitally important^10^. As of 2024, at least 42 CMS-related missense variants have been reported to affect AChR kinetics^7,11–20^.

While protein structures are often key to understanding protein function or the effects of pathogenic mutations, structural study of the muscle-type AChR is particularly challenging due to difficulties in the recombinant expression of hetero-multimeric channels. Accordingly, existing receptor structures are of orthologs purified from native tissues. Receptor structures from the *Torpedo* electric ray have served as prototypes for AChR since the first 9 Å reconstruction in 1993, revealing its overall architecture, as well as its modulation by ligands, anaesthetics and toxins^21–27^. More recently, bovine AChR structures shed light on structural components contributing to the different electrophysiological signatures of its foetal and adult isoforms^28–35^. Despite these advances, the structure of human muscle-type AChR remains elusive. Critically, the sequence of bovine AChR is only 92% identical to the human receptor, this difference may affect interpretation of AChR variants in relation to diseases such as CMS.

To address this ambiguity in the structure and function of human AChR, we resolved the structures of the human muscle-type receptor (α_2_βδε) in α-bungarotoxin-(αBuTx-) bound resting states and acetylcholine- (ACh-) bound desensitised states using cryo-electron microscopy (cryo-EM). By comparing the two states of AChR, we uncovered structural changes associated with its recovery from desensitisation. In addition, we used single-channel electrophysiology and structural models to understand the molecular consequences of eight CMS-causing AChR variants.

## Results

### Structure of the resting state human α_2_βδε AChR

In order to assemble an adult muscle-type AChR for structural and functional analyses, we generated a doxycycline-inducible cell line that simultaneously expresses human wildtype (WT) α, β, δ subunits along with an ε subunit containing an enhanced green fluorescent protein (EGFP) insertion into the M3-M4 intracellular loop to monitor protein expression. After confirming the receptor is properly folded and assembled, we obtained cell-attached single channel recordings of this stable cell line which showed WT AChR-like conductance and kinetics (Figure 1A-1B), consistent with previous recordings of the transient expression of the same construct^36^ and functional neuromuscular junctions in transgenic mice^37^. Recombinant AChR was solubilised in a detergent mixture of 1% n-dodecyl-β-D-maltopyranoside (DDM) and 0.1% cholesterol hemisuccinate, and then purified to homogeneity using both affinity and size-exclusion chromatography (Figure S1, Table S1). For cryo-EM, we added Fab35^38^, an anti-α IgG1 fragment, to act as a fiducial label and break the receptor’s 5-fold pseudosymmetry.

**Figure 1:**
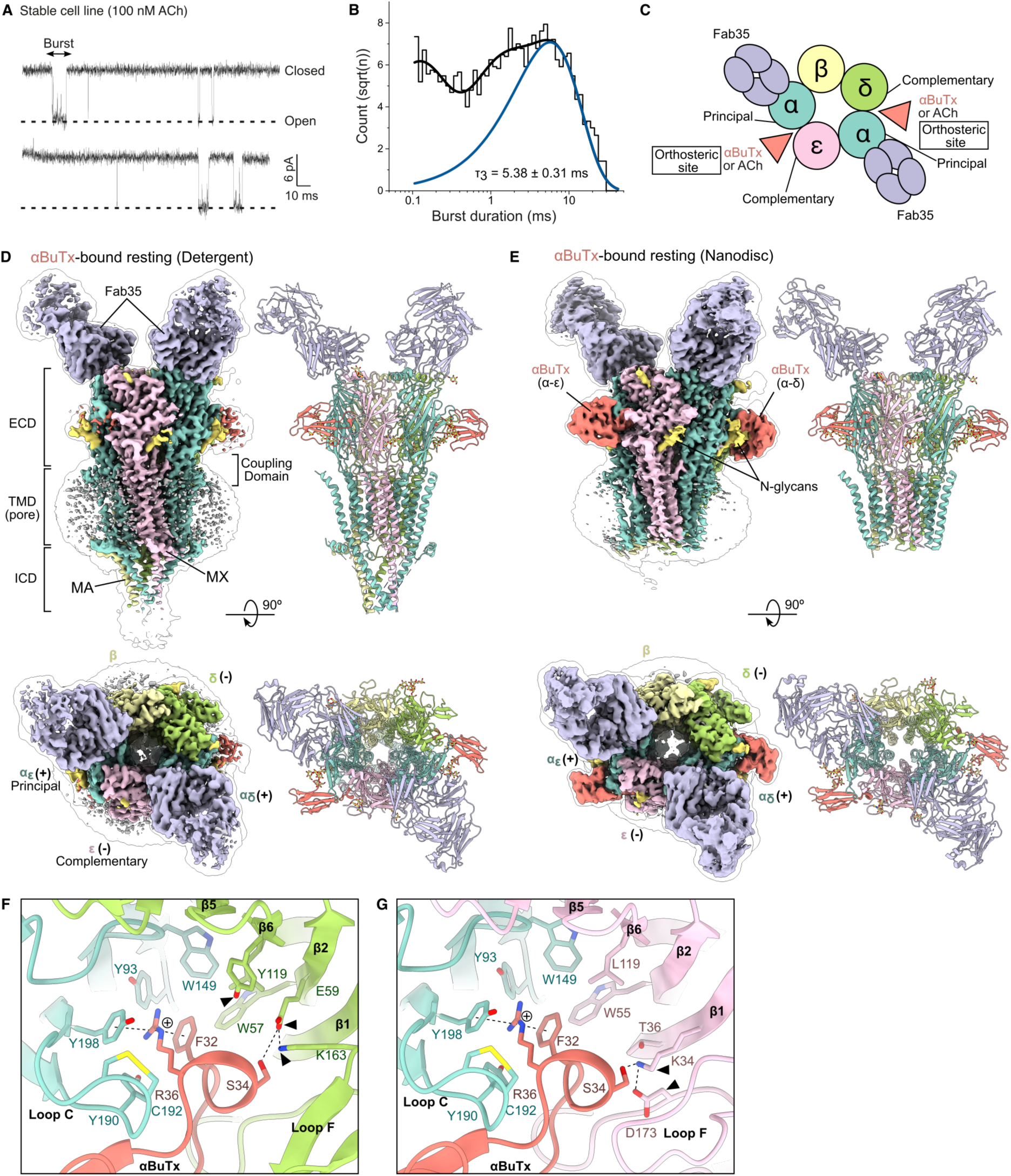
AChR in the αBuTx-bound resting state. (A-B) Single-channel currents and burst duration histograms produced by AChR-expressing stable cell line at +80 mV in the presence of 100 nM ACh, τ_3_=5.38±0.31 ms, n=4. (C) Schematic of AChR stoichiometry when viewed from the synapse. Orthosteric sites are formed by the principal α subunit and the complementary δ/ε subunit. (D-E) Cryo-EM structures of the detergent- and nanodisc-reconstituted αBuTx-bound receptor. (F-G) Comparison of αBuTx binding to the α-δ and α-ε interface in the nanodisc structure. Black triangles indicate key non-conserved residues in δ/ε subunits; cation-π interactions are denoted by dotted lines. ECD, extracellular domain, TMD, transmembrane domain, ICD, intracellular domain.

To obtain a resting-like structure of AChR, purified protein was incubated before vitrification on cryo-EM grids with αBuTx, an 8 kDa α-neurotoxin from *Bungarus multicinctus* venom that prevents channel activation^39,40^ While AChR pentamers bound to either two Fabs or one Fab were present during cryo-EM data processing (Figure S2), likely a result of complex dissociation at the air-water interface during sample vitrification, we could nevertheless observe well-resolved structural features and side chains within these maps. To better resolve the mobile transmembrane domain (TMD), the two Fab subset of particles were futher classified based on TMD conformation (Figure S2). Ultimately, despite the significant compositional and structural heterogeneity present, we were able to select a subset of particles that yielded a 2.96 Å reconstruction (Table S2). The protein was well-resolved throughout its extracellular (ECD), TMD and intracellular domains (ICD). In general, the map quality enabled modelling of the protein into the experimental map using N-glycans and identifiable sequence motifs to identify each chain, numbering each modeled chain after excluding the signal peptide, confirming we had captured the expected stoichiometry of α_2_βδε.

The human AChR structure retain the classical architecture described for the previously reported *Torpedo* and bovine structures^21–28^ (Figure S3A-E). The receptor subunits are arranged in the clockwise order α-β-δ-α-ε when viewed from the synaptic cleft (Figure 1C-D), with one Fab35 bound to the N-terminus of each α subunit. The αBuTx binding sites are at the α-ε and α-δ extracellular interface which coincides with the two orthosteric sites (Figure 1C-D). The extracellular domain of each subunit consists of a β-sandwich (β1-β10), followed by four membrane-spanning helices (M1-M4), with the M2 helix lining the pore. The C-terminus of M3 helix is followed by the MX helix that lies parallel to the inner leaflet of plasma membrane. The MX helix then transitions into a disordered intracellular loop that becomes helical once more at the intracellular MA helix precedes the M4 helix. We refer to the unresolved region between M3 and M4 helices as the M3-M4 loop.

Surprisingly, we observed poor occupancy of αBuTx in the Coulomb potential map, with the α-δ interface being the higher occupancy site (Figure S4A). We hypothesise this partial occupancy of αBuTx was due to reduced affinity of the detergent-solubilised receptor for toxin. We also noted weak TMD density, corresponding to mobility in that domain, likely due to detergent molecules destabilising the interhelical interactions in the TMD^41^. As these shortcomings in the detergent-solubilised AChR map make analysing these regions difficult, we reconstituted the purified AChR into MSP2N2 lipidic nanodiscs to create a more native-like environment and collected a second cryo-EM dataset. We isolated particles of AChR in nanodiscs which produced a 2.48 Å reconstruction map, with 2 Fabs bound and clear density for the distinctively shaped long C-terminal tail on the δ subunit that is similar to structural homologs (Figure S5, Table S2)^24^. In this structure, both αBuTx binding sites were fully occupied, but the receptor’s TMD and ICD appeared more dynamic than the detergent-solubilised structure, with no resolvable density for the ICD’s MX and MA helices (Figure S6A-C, Figure 1E).

In our αBuTx-bound structures, residues αBuTx-R36 and αBuTx-Y32 extend into the aromatic cage of the binding site, lined by αW149, αY190, αY198 and δW57 or εW55 (Figure 1F). We noted that the αBuTx binding mode to the human receptor is similar to that of the *Torpedo* receptor, consistent with having highly conserved interacting residues (Figure S3F-H)^42,43^. The αBuTx competes with ACh and stabilises the open conformation of α loop C, thus trapping the receptor in the resting state. The guanidinium group of αBuTx-R36 is sandwiched between αBuTx-F32 of the toxin and αY198 in a stacked cation-π interaction. Comparing the δ and ε subunits, we noted residue differences between the two ligand binding sites. On the δ subunit, δE59 on β2 forms a salt bridge with δK163 on loop F, while δE59 is also hydrogen bonded to αBuTx-S34. The equivalent salt bridge is reversed on the ε subunit, with εK34 of β1 and εD173 of loop F, whilst its hydrogen bond contact with αBuTx-S34 is maintained by εK34 (Figure 1F-1G). ^42,43^

### Structural changes between desensitised and resting states

We next sought to characterise the structural changes between desensitised and resting states by determining acetylcholine-bound AChR. Samples were prepared in nanodisc as before, replacing αBuTx in the sample buffer with either 1 mM ACh or a triad of 100 μM ACh, positive allosteric modulator (PAM), and fluoxetine (see Methods). The 100 μM ACh dataset gave rise to a 2.73 Å reconstruction, whilst the 1 mM ACh dataset produced a 2.64 Å map (Figures S7-S8, Table S2). The 1 mM ACh dataset suffered from severe preferential orientation of the particles resulting in poor resolution of the TMD (Figure S6D).

Consequently, we analysed the 100 μM ACh-PAM-Fluoxetine structure to analyse the desensitised state (Figure 2A), as both structures were bound to ACh only (Figure S4B) and adopted identical desensitised conformations (RMSD= 0.856 Å; Figure S3B, Figure S9A), which we atributed to being a desensitised conformation because of a high degree of similarity with published desensitised structures (Figure S3C-E). Our observed weak TMD density may also be caused by the previously noted collapsed pore phenomenon for purified heteromeric pLGICs^44,45^, which may be caused by suboptimal composition of the membrane mimetic^46–49^.

**Figure 2:**
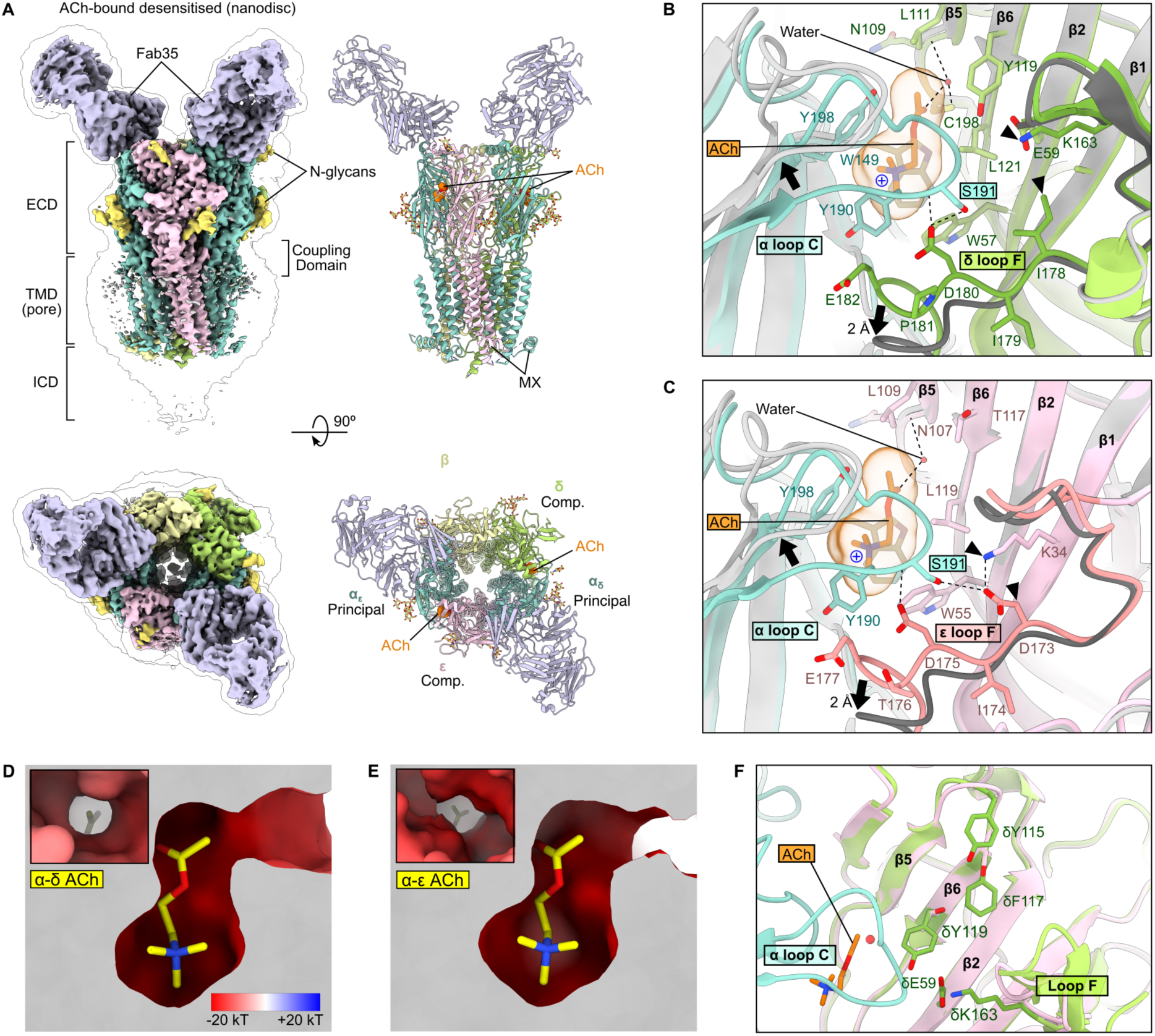
ACh binding sites in the desensitised state. (A) Cryo-EM structure of the nanodisc-reconstituted ACh-bound receptor. (B-C) ACh binding site is stabilised by hydrogen bonding between αS191 and δ/ε loop F. For comparison, resting state structure in detergent is shown in grey, overall RMSD=0.878 Å. Black triangles indicate differences between δ and ε subunits, black arrows indicate direction of movement associated with structural changes from desensitised to resting state. (D-E) ACh binding site is more occluded and less electronegative at the α-δ site compared to the α-ε site. (F) Bulky residues (δY115, δF117, δY119, δE59 and δK163) within 15 Å of the α-δ ACh binding site, equivalent ε residues are εG113, εS115, εT117, εG57 and εA161).

The ACh binding sites are formed by α loop C on the principal α subunit, and β strands β2, β5 and β6 on the complementary δ/ε subunits, with the pocket enclosed by the interaction between α loop C and δ/ε loop F (Figure 2B-C). Compared to the αBuTx-bound structure, ACh binding is accompanied by the complete closure of α loop C, and this action is blocked by αBuTx binding. The positively charged tertiary amine group of ACh is stabilised by cation-π interactions with αY190, αW149 and αY198 on the principal subunit, and with εW55 or δW57 on the complementary subunit. The acetyl group of ACh is coordinated by a water at both α-δ and α-ε orthosteric sites. These structural waters mediate an interaction between the carbonyl group of ACh and the mainchain carbonyl of εN107. Notably, the pose of ACh is slightly different to that of a bovine AChR structure, due to an intact disulphide between αC192 and αC193 in our human AChR structure, whereas the equivalent bond is broken in the bovine receptor structure^28^ (Figure S3I-J).

The closed conformation of α loop C in the ACh-bound state is stabilised by a hydrogen bond network between αS191 on loop C and acidic residues on δ/ε loop F, a role that was previously described in AChR ortholog structures to stabilise the closed conformation of the α loop C (Figure 2B-C)^26,28,50^. Notably, the ε subunit contains one more acidic residue than the δ subunit, with this charge introduced by εD173. The ACh binding cavity appeared less occluded and more electronegative at the α-ε interface (Figure 2D-E). The occlusion at α-δ site may be attributed to the five bulky residues δY115, δF117, δY119, δE59 and δK163 within 13 Å of the bound ACh at the α-δ site, while the equivalent residues εG113, εS115, εT117, εG57 and εA161 on the ε subunit are smaller (Figure 2F). The bovine AChR contains the same key residues at the α-δ and α-ε interfaces except for a serine to alanine substitution at αS191, resulting in the loss of hydrogen bonding between α loop C and δ/ε loop F.

Our aBuTx and ACh-bound receptor structures show differences beyond the orthosteric site, revealing how local changes at this site transmit to the TMD via the coupling domain. In the desensitised structure, we identified a key hydrogen bond interaction between αS266 of the M2-M3 loop and εE184/δE189 of loop F (Figure 3A-C)^51^. The acidic loop F residues are also involved in a conserved tripartite salt bridge of εR218, εE184, εD45, and εD138 at the base of ECD (Figure 3B-C). Interestingly, the αS266-εE184 hydrogen bond is observed only in the desensitised state, but the αS266-δE189 interaction persists in both resting and desensitised states. We hypothesise the structural transition from desensitised state back to resting state involves first the dissociation of ACh, followed by the concerted opening of α loop C and a 2 Å downward movement of δ/ε loop F, uncoupling the hydrogen bond between αS191 and loop F of the δ or ε subunits (Figure 2B-C, dotted lines). These conformational changes are transmitted down to the coupling domain, which leads to αS266 of the M2-M3 loop moving away from the δE189/εE184 also on loop F (Figure 3B-C, yellow arrows). Moreover, the α_δ_ M2-M3 loop undergoes a 2 Å shift at αS266 and a 4 Å shift for the same residue on α_ε_, thus contributing to the asymmetrical conformational changes within the AChR pentamer. Ultimately, M2-M3 loops propagate the signal of ACh binding to the TMD, where the alignment of five M2 helices determine the channel’s ion permeability.

**Figure 3:**
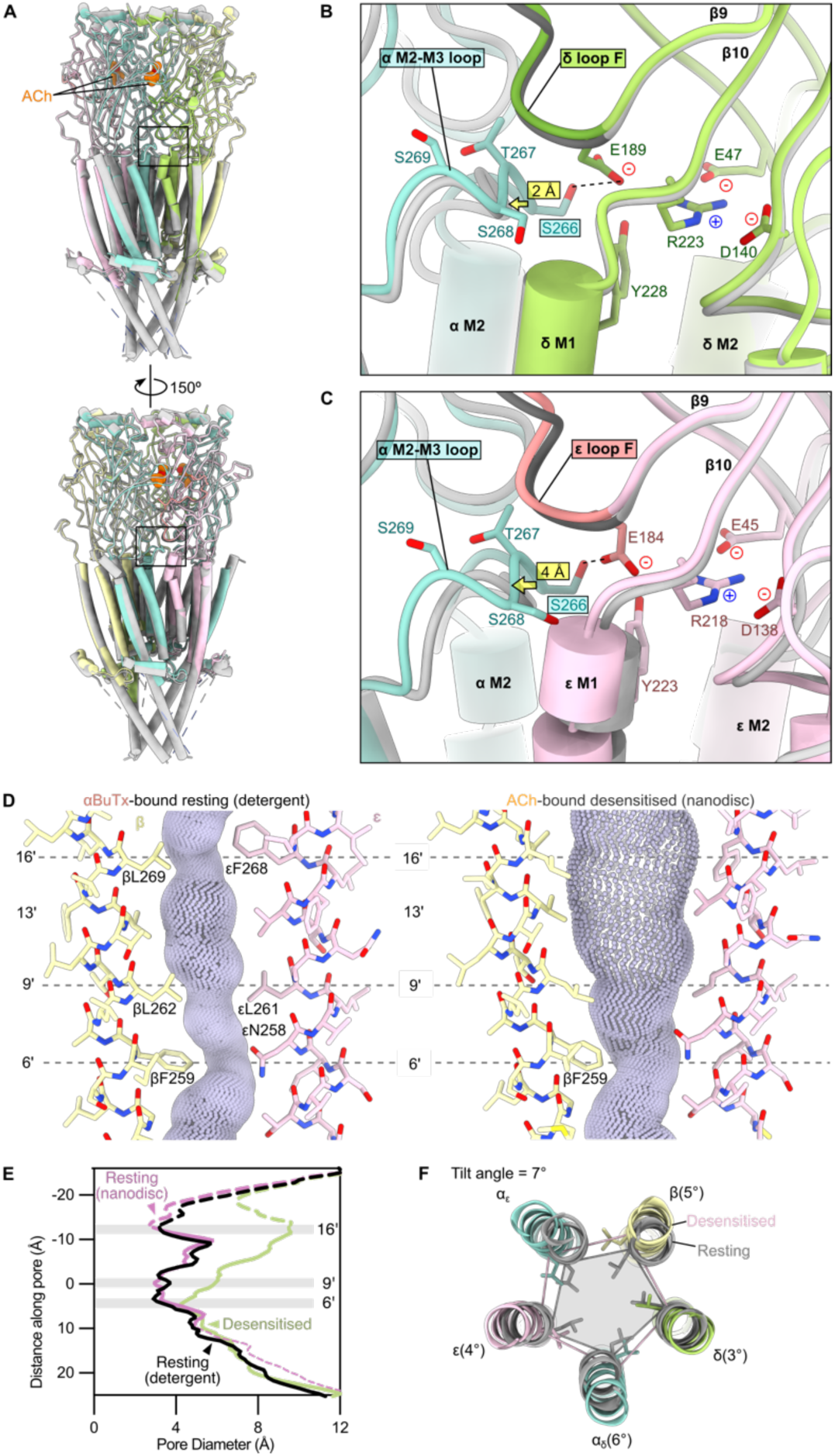
Structural transitions between desensitised and resting states. (A) Structural overlay of desensitised (coloured) and resting (grey) states, overall RMSD=0.859 Å. Boxes highlight the positions of inter-subunit interactions in the coupling domain. (B) Hydrogen bonding between αS266 and δE189 as part of a tripartite salt bridge on the complementary subunit (δR223-E189-E47-D140). (C) The same interaction between αS266 and εE184 at the equivalent position. (D-E) Comparison of pore profiles of resting and desensitised states. Dashed lines in (E) indicate regions where some sidechain positions could not be experimentally determined but were reinstated for the purposes of profile generation (see Methods). (F) Pore dilation is achieved by 9’ leucine residues rotating away from central axis, accompanied by outward tilting of the M2 helices: 7° (α_ε_), 5° (β), 3° (δ), 6° (α_δ_) and 4° (ε).

By convention, residues in the M2 helices of pLGICs are numbered from -1’ on the intracellular side, equivalent to αE241, to 20’ on the extracellular side, equivalent to αE262. For the αBuTx-bound structure, its pore is narrow throughout, which contains a 2.9 Å constriction at the 9’ hydrophobic gate (Figure 3D). In contrast, the desensitised ACh-bound channel has a funnel-shaped pore, wider at the 9’ position with a pore diameter of 6 Å, which tapers to a 4.2 Å constriction at the 6’ position (Figure 3D-E). Both pore conformations are similar to the corresponding resting and desensitised states of bovine and *Torpedo* receptors^24,26,28^ (Figure S9). In our structures of the desensitised human AChR, the wider 9’ gate results from the five leucine residues rotating away from the pore axis (Figure 3F). At the 6’ position, βF259 causes a visible kink in the permeation pathway in all three structures with well-resolved TMDs, forming the desensitisation gate that excludes the passage of hydrated Na^+^ ions. This finding is supported by the observed faster desensitisation of a βF259S targeted mutant^28^. Apart from the rotation, the M2 helices also tilt away from the pore axis in the desensitised state, with tilt angles ranging from 3° to 7° (Figure 3F, Figure S9I)^28^.

### Characterisation of eight congenital myasthenic syndrome variants

Using our human AChR structures, we next sought to understand the effects of eight previously unstudied CMS AChR kinetic variants gathered from many years of clinical surveillance (Table S3, Figure 4A). The variants were found to have negligible effect on the surface expression of AChR, defined as ≥50% of WT, indicating that altered AChR kinetics rather than AChR deficiency likely underlies these patients’ syndrome (Figure 4B). Therefore, we used single-channel recordings and burst duration statistics of the variants to classify the dominant pathogenic mechanism as either FCCMS or SCCMS, thereby enabling tailored therapeutic intervention. Simply defined, FCCMS is characterised by shortened burst duration compared to WT, whilst SCCMS is characterised by prolonged bursts (Figure 4C). Three variants, αT133I, βI285S, and δD180N, led to a reduction in the longest time constant of AChR burst duration (1.86±0.13, 1.81±0.29 and 1.05±0.04 ms) compared to WT (5.61±0.38 ms) (Figure 4D-E, Table S4). The remaining five substitutions severely prolong the AChR bursts, with time constants ranging from 18.11±2.43 ms for αT281S to 69.88±0.07 ms for εS235A (Figure 4D-E, Table S4). Whilst most of these CMS variants did not alter the single channel conductance of AChR, the αI49T slow-channel variant resulted in markedly decreased current amplitudes at different holding potentials, with a single channel conductance of 33 pS compared to 63 pS of WT receptors (Figure 5A-B).

**Figure 4:**
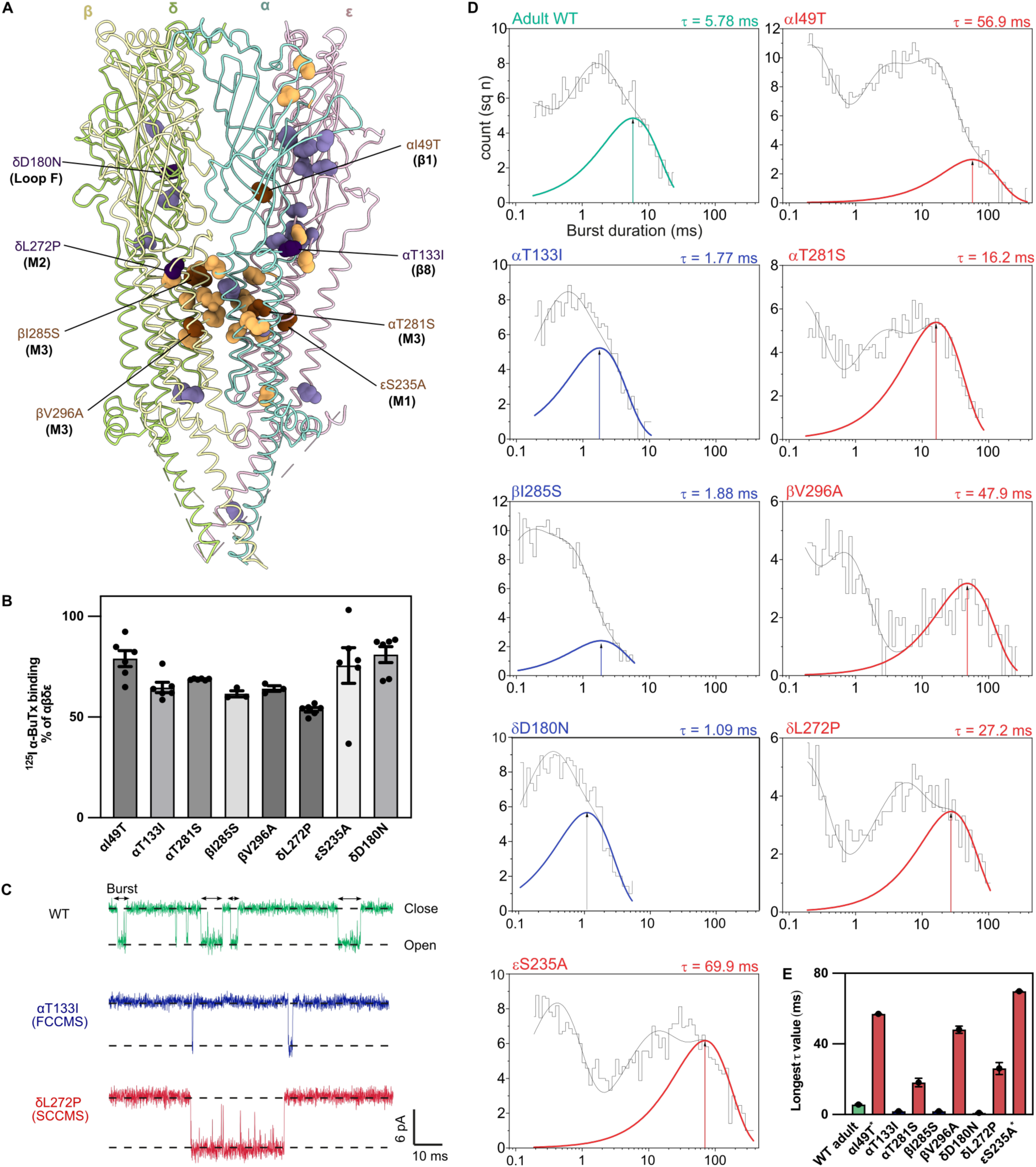
Characterisation of CMS kinetic variants. (A) Locations of SCCMS (brown/orange spheres) and FCCMS variants (purple/lilac spheres) mapped onto the detergent-reconstituted resting state structure^7,11–19^. Variants reported in this study are highlighted in a darker shade and labelled, α_δ_ subunit is omitted for clarity. See Table S5 for the full list of CMS variants on the diagram. (B) Surface expression of CMS variants in transfected HEK293 cells measured by ^125^I-αBuTx binding assay. Measurements were normalised to the expression of WT receptors, n≥3. (C) Representative single channel currents of WT, fast channel (αT281I), and slow channel (δL272P) variants at +80 mV holding potential in the presence of 100 nM extracellular ACh, n≥4. Channel openings are upward deflections. (D) Burst duration histograms fitted with sum of exponentials for WT, slow channel variants (αI49T, αT281S, βV296A, δL272P, and εS235A) in red, and fast channel variants (αT133I, βI285S, and δD180N) in blue. The δD180N variant was recorded with 500 nM instead of 100 nM ACh. (E) Summary bar chart of the longest burst duration time constant (τ) extracted from the histograms. Data is presented as mean ± SEM, except for entries indicated with asterisks (*) which had fitting error instead of SEM (see Table S4).

**Figure 5:**
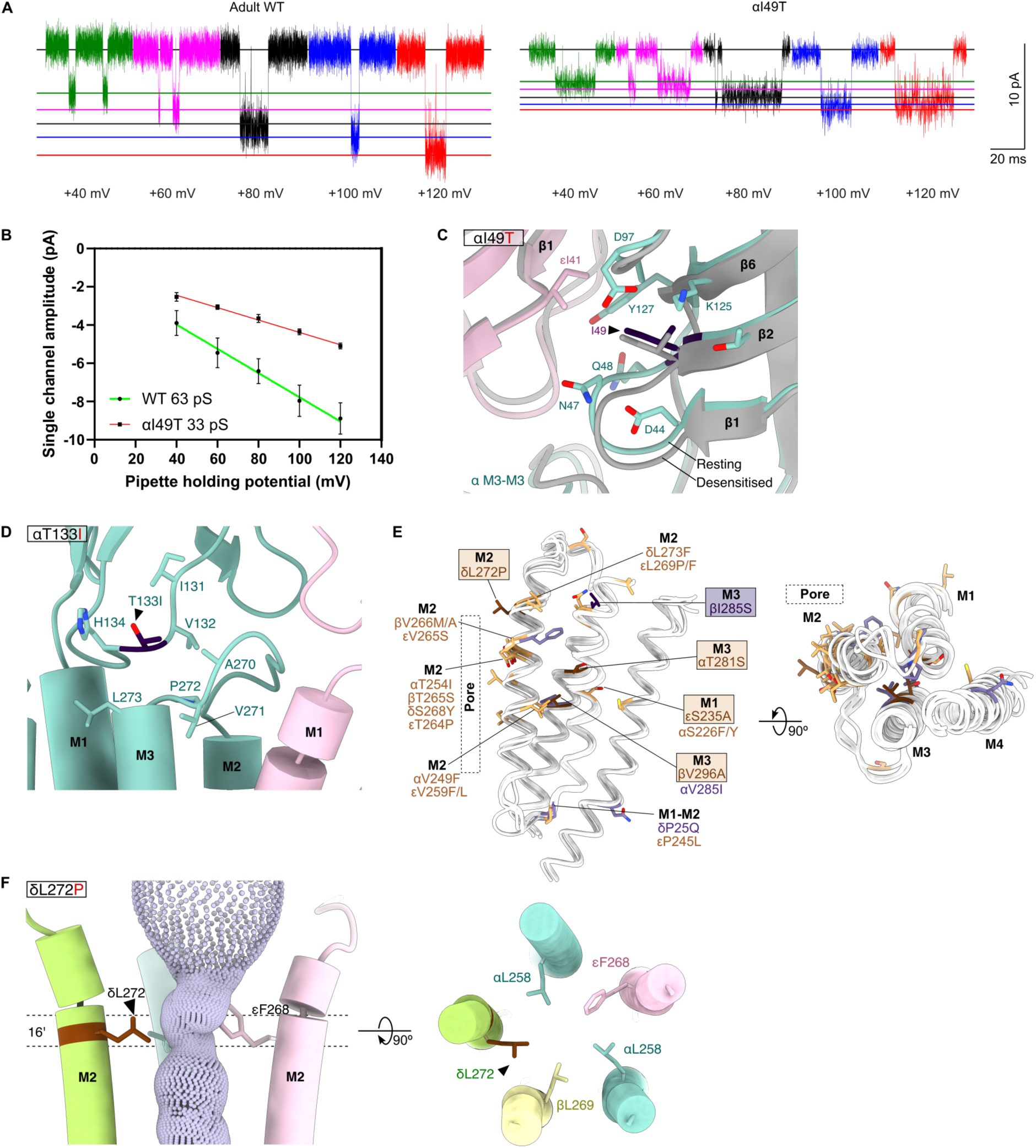
Structural analysis of CMS kinetic variants. (A-B) αI49T reduces AChR single channel conductance from 63±11 pS (WT) to 33±3 pS. Current amplitudes of bursts recorded at +40 to +120 mV were plotted in a current-voltage curve, and single channel conductance was the slope of its linear regression. (C) αI49T at the extracellular vestibule does not undergo large conformational changes between resting and desensitised (grey) states. (D) αT133I is located at the coupling domain. (E) Mapping of CMS kinetic variants found in the transmembrane domain, FCCMS in purple, SCCMS in orange. Variants are shown on superposed TMDs. (F) δL272P replaces a side chain at the 16’ pore constriction.

Examining our newly determined structures of human AChR to better understand the structural consequences of these variants, three CMS variants are located within the extracellular domain of AChR (αI49T, δD180N and αT133I). The majority of SCCMS variants are located in the AChR TMD, whereas FCCMS variants may be scattered across all domains^7^ (Figure 4A, Table S5). The αI49T variant, which causes SCCMS and a reduced conductance, is found in the extracellular vestibule and 11 Å from M2-M3 of the coupling domain (Figure 5C). Though its surroundings remain mostly static between desensitised and resting states, the introduction of a threonine increases the polarity of this region in the ion permeation pathway, though ultimately how this causes such drastic changes in channel kinetics and conductance is unclear. Another fast channel variant, δD180N altered the charge of the key δD180 loop F residue responsible for stabilising α loop C in the ACh-bound state (Figure 2B), providing a structural explanation to its loss-of-function characteristics. A previously identified variant at the equivalent εD175N position also causes FCCMS^52^. The third FCCMS variant, αT133I, is in the Cys-loop and immediately adjacent to the previously reported αV132L which also causes fast channel syndrome^53^ (Figure 5D). Both αT133I and αV132L appear to exert their effects by increasing the hydrophobicity of the Cys-loop which may affect its coupling with the M2-M3 loop.

The remaining five CMS variants are located within the transmembrane domain, εS235A in M1, δL272P in M2 and βI285S, βV296A and δL272P in M3 (Figure 5E). Firstly, the δL272P slow channel variant removes a key side chain at the 16’ constriction of the pore, whilst possibly disrupting the helicity within M2 through the introduction of more conformationally restricted amino acid (Figure 5F). This pore-facing side of the M2 helix is a slow channel hotspot (Figure 5E), with the SCCMS variants αT254I, βT265S, δS268Y, and εT264P at the 13’ position which lies one-helix turn below δL272P. Similarly, βL262M is located a further helix-turn away at the 9’ hydrophobic gate^7^. This suggests missense variants in M2 likely affect channel gating by altering its ability to rotate open or shut in response to ACh binding.

Similar to the M2 SCCMS variants clustering on that helix, we noted the εS235A variant on M1 coincides with the αS226F and αS226Y SCCMS variants on the α subunit^7^. Furthermore, of the βI285S, βV296A and αT281S FCCMS variants on M3 described here, βV296A is located at the equivalent position of the αV285I FCCMS variant on the α subunit^54^. Notably, most M1 and M3 kinetic variants face the hydrophobic core of the four transmembrane helix bundles and are likely to affect proper helix packing and alignment.

### N-terminus of the α subunit contains a metal-binding motif

In examining our AChR structures determined on copper grids, αBuTx-bound in nanodisc and 100 μM ACh in nanodisc, we observed strong non-protein density at the N-terminus of the α_δ_ subunit (Figure 6A-B, Figure S10A). Its residues 1-3 adopt a square planar arrangement, with their backbone nitrogen and a fourth nitrogen from the H3 imidazole group coordinating an unknown ion, which presumably is a Cu^2+^ derived from the EM grid (Figure 6B, Figure S10). In contrast, the same residues in the detergent-solubilised αBuTx-bound sample on gold grids exhibits an open conformation (Figure 6C). This was immediately reminiscent of the Amino Terminal Copper and Nickel (ATCUN) sequence motif X1-X2-H3, which was first described to coordinate divalent cations in human serum albumin^55^. This sequence motif is found only on the muscle-specific α1 subunit as S1-E2-H3, after removal of signal peptide, and it is conserved in both bovine and *Torpedo* orthologs (Figure 6D). Based on the location of the ATCUN sites within the structure, metal binding may be influenced by Fab35 binding, as well as the neighbouring ε and β subunits (Figure 6E, Figure S10).

**Figure 6:**
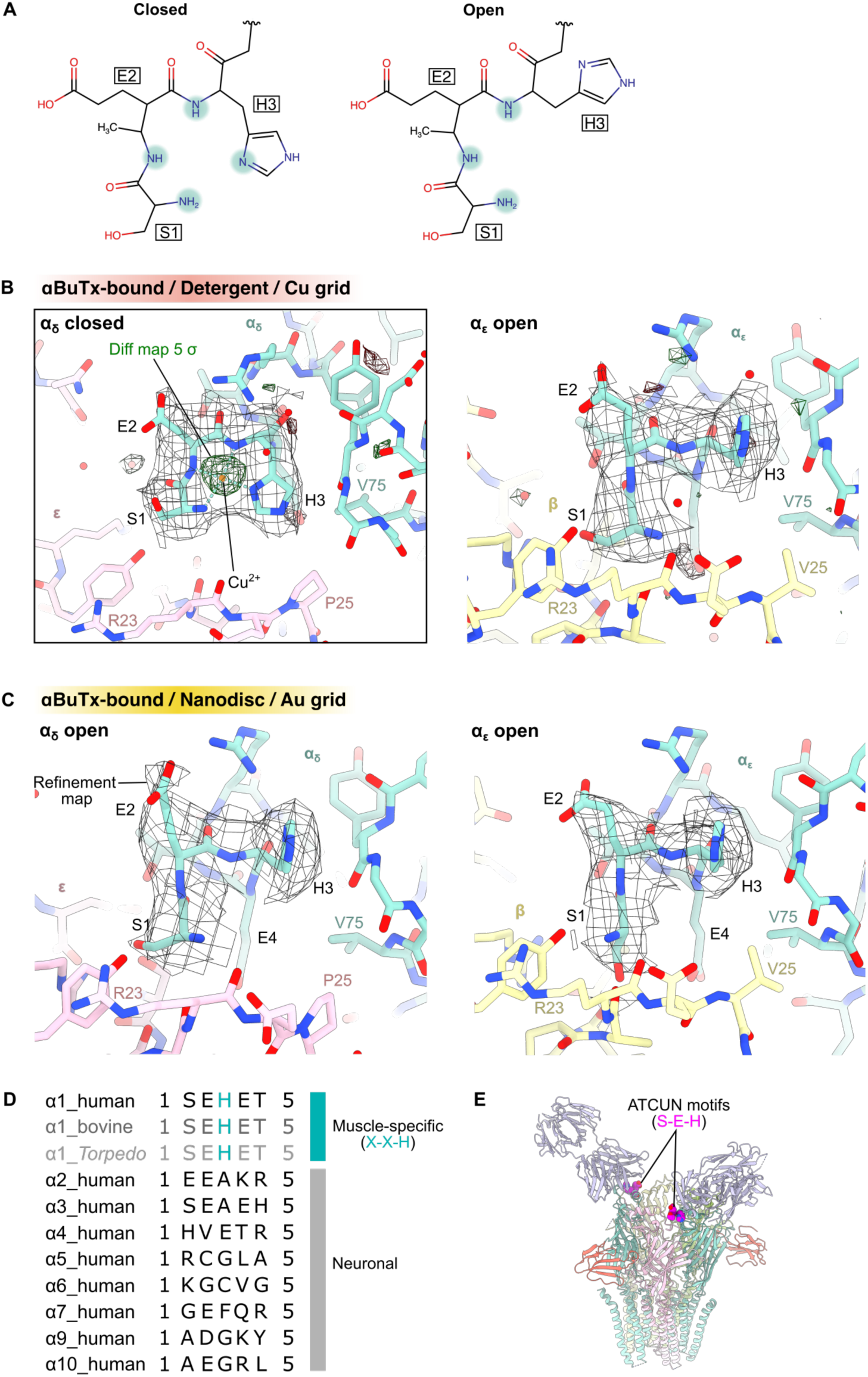
N-terminus of the α subunit contains a metal-binding motif. (A) Schematic of the ‘X1-X2-H3’ amino terminal Cu^2+^ and Ni^2+^ (ATCUN) motif in ion-coordinating (closed) and open conformations. (B-C) Comparison of ATCUN conformations at the α_δ_-ε and α_ε_-β interfaces in structures solved from copper (in nanodisc) versus gold (in detergent) cryo-EM grids. Coulomb potential maps are shown in grey, mask-normalised F_o_-F_c_ difference maps highlighting non-protein features are shown in green (positive density) and red (negative density), calculated with Servalcat (see Methods). (D) Comparison of N-terminal sequence of AChR α subunits. Sequences were sourced from UniProt^56^ with signal peptides omitted. (E) ATCUN motifs are located at the N-terminus of the α subunits near Fab35 binding sites.

### Recombinantly-expressed AChR can assemble into multiple stoichiometries

When processing the cryo-EM dataset of detergent-solubilised AChR, we isolated channels with subunit compositions other than the expected α_2_βδε, indicating that multiple hetero-pentamer species were expressed in the recombinant cell line. To identify the oligomers present in our sample, we used ModelAngelo^57^, to automatically identify and build the various well-resolved classes, and the results were confirmed by visual inspection of regions with distinctive protein sidechains (Figure 7A-B). We identified particle subclasses with α_3_ε_2_ (Figure 7C-D), where tilting of the ε M4 and a poorly resolved TMD suggest this receptor has a collapsed pore and is likely non-functional. We also identified a tentatively assigned α_2_βδ_2_ subclass with a well-resolved TMD and ICD (Figure 7E-F). To probe whether this α_2_βδ_2_ receptor is functional on the cell surface, we transfected HEK293 cells with cDNAs encoding only the α, β, and δ subunits and observed ACh-elicited single channel currents with current amplitude between that of adult WT and foetal WT receptors (Figure 7G). This demonstrates that functional receptors can form on the cell surface without the ε/γ subunit. However, both automated subunit assignment using ModelAngelo and visual analysis indicated sequence ambiguities in its second δ subunit (in place of ε), which suggests a mixed δ/ε population at this subunit position (Figure 7F-G). Further classification was unable to resolve a complete ‘clean’ α_2_βδ_2_. Nevertheless, this suggests although never reported, other AChR pentamers may be present at the human neuromuscular junction, or alternately, receptor assembly at the NMJ is regulated by proteins absent in our HEK293-based expression system.

**Figure 7:**
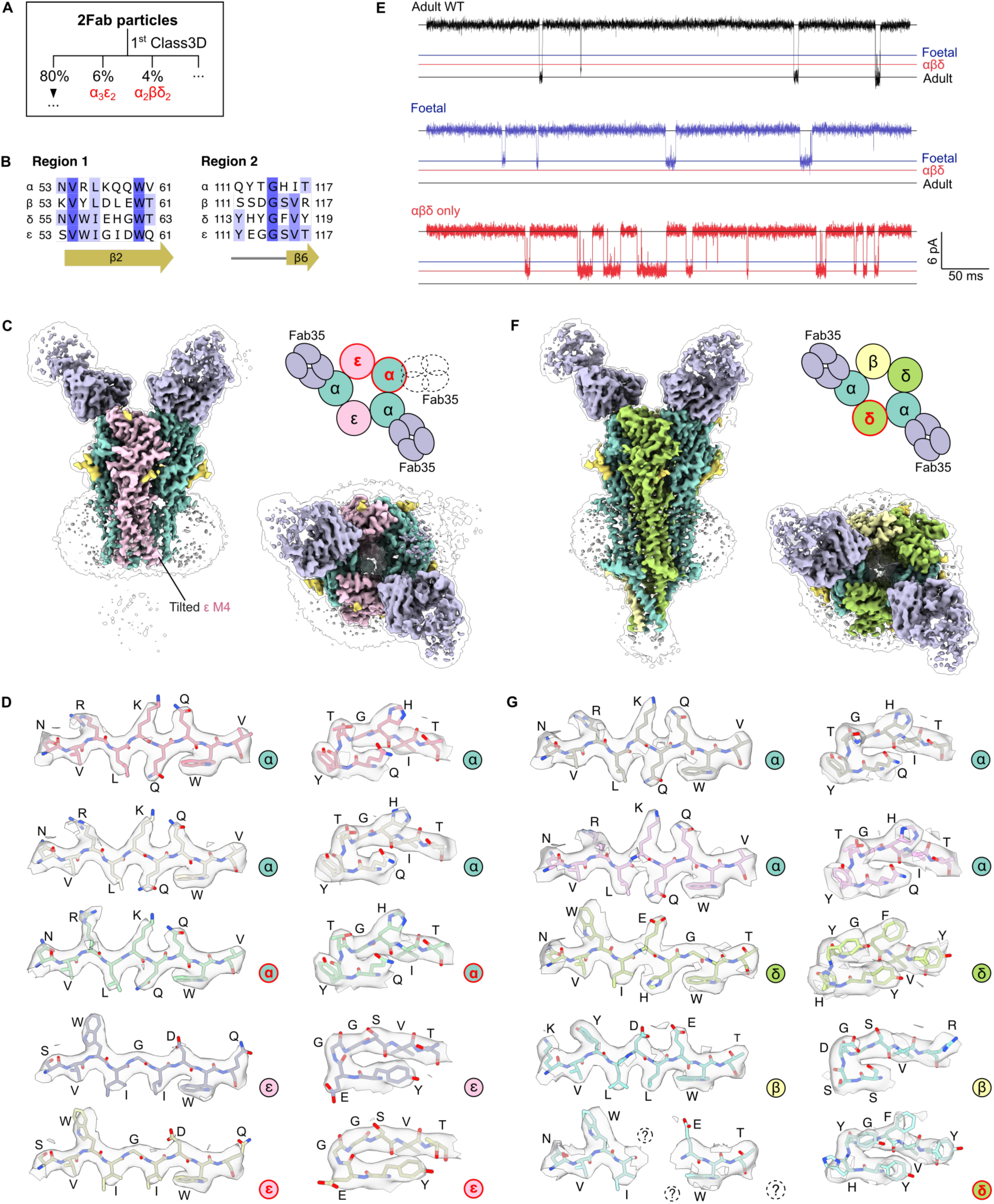
Recombinantly-expressed AChR can assemble into multiple stoichiometries. (A) The α_3_ε_2_ and α_2_βδ_2_ particles were isolated from the 2Fab particles during RELION^58^ Class3D of the detergent-solubilised cryo-EM dataset. (B) Two ECD regions (equivalent to α53-61 and α111-117) with distinctive protein sidechains were used for visual confirmation of subunit assignment. Structural models for α_3_ε_2_ and α_2_βδ_2_ particle subclasses were built by ModelAngelo based on protein sequence and Coulomb potential maps^27^. (C) A 3.2 Å map of α_3_ε_2_ was reconstructed from 25,453 particles. (D) Model-map agreement for α_3_ε_2_ subunits in regions 1 and 2. (E) Single channel currents from HEK293T cells expressing adult WT (αβδε), foetal WT (αβδγ), or αβδ subunits at +80 mV with 100 nM ACh. (F) A 3.2 Å map of predominantly α_2_βδ_2_ as reconstructed from 18,001 particles. (G) Model-map agreement for α_2_βδ_2_ subunits in regions 1 and 2.

### Structural determinants of adult versus foetal AChR

The muscle-type AChR transitions between a foetal receptor to an adult receptor during development^59^. As both adult and foetal receptors consist of the same α, β and δ subunits, functional differences, such as adult receptor’s shorter burst duration and higher conductance, are mostly down to the sequence differences between ε versus γ subunits which are 48% non-identical (Figure 8A in blue, Figure S11). Previous studies suggested that residues in the β5-β6 hairpin of ε ECD dictate the adult receptor’s open probability^60^, and acidic residues in the ε ICD contribute to its higher channel conductance^28^. We set out to investigate other regions of notable structural differences between the foetal and adult receptors. We used the structure of bovine γ subunit (PDB: 9AWK) to guide this analysis as it has an identical sequence to human AChR in the loop F and M2-M3 loop^28^ (Figure S11). Examining our structures of the adult isoform AChR, we identified three ε-specific motifs that warrant further investigation: εK34-εD173 salt bridge, a εC190-εC470 disulphide bond, and a εS280 M2-M3 loop residue that interacts with εT133 of Cys-loop (Figure 8A-C). Firstly, we noted an εK34-εD173 salt bridge and an analogous interaction at δE59-δK163 earlier (Figure 1F-G). This salt bridge is replaced by γK34-γE163 in the foetal subunit, with an equivalent lysine residue as εK34 but a different acidic residue (Figure 8A). To test the importance of the εK34-εD173 interaction, we introduced the γ acidic residue by the εD163E substitution, and in a separate experiment, combined it with a second εD173F mutation to knock-out the ε acidic residue. The εD173F mutation severely reduced surface expression, whereas the double mutant has 60±2% of the surface expression of WT receptors (Figure 8D) and it did not alter the receptor’s burst duration (Figure 8E, Table S4). Secondly, an ε-specific εC190-εC470 disulphide bond links its C-terminus to the ECD (Figure 8A), and the same bond is observed on an ε-containing bovine AChR structure^11^. We found the εC190A substitution severely reduced receptor expression, which corroborated a previous study that probed the other cysteine residue via a εC470A mutation^26^. Thirdly, we focussed on a M2-M3 loop residue, εS280, which interacts with εT133 of the Cys-loop (Figure 8C). Substituting εS280 to an alanine, as present in the γ subunit, caused a marked reduction in AChR surface expression (Figure 8D). We conclude that all three sites on the ε subunit are key to receptor expression, though due to the low expression, we were unable to assess the effects on channel kinetics of εD173F, εC190A and εS280A.

**Figure 8:**
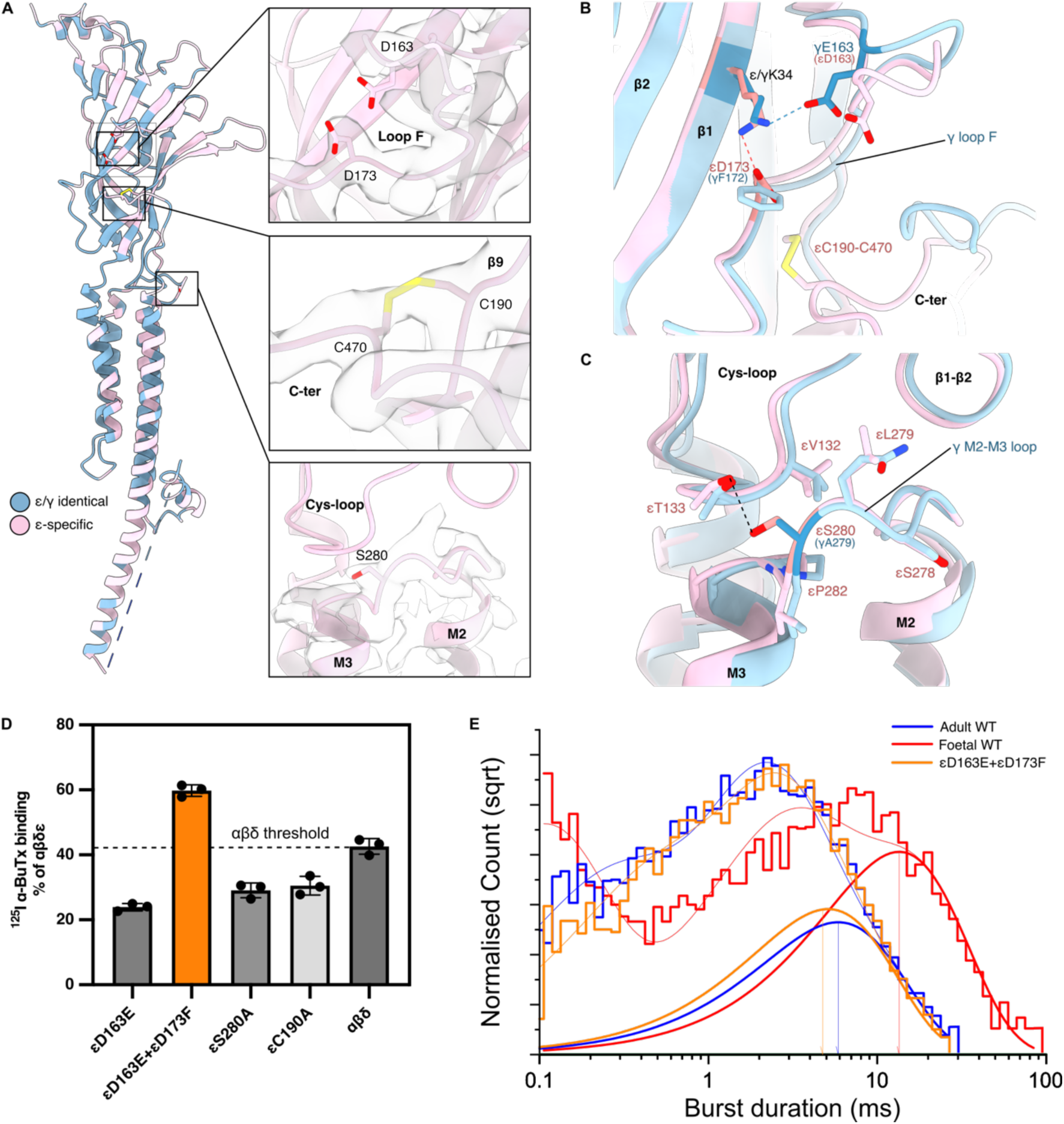
Characterisation of ε to γ mutants. (A) Structure of the resting state ε subunit coloured by ε-specific residues (pink) or ε/γ consensus residues. Positions of mutated ε residues on loop F (D163 and D173), β9 strand (C190), and M2-M3 loop (S280) shown on the structure. The resting state human ε subunit adopts the same conformation as the resting state bovine γ subunit (PDB: 9AWK), RMSD=0.658^28^. (B) εD173F forms a salt bridge with εK34 in the resting state. εD163E knocks in the equivalent γ residue, and εD163E+εD173F knocks out the formation of ε-specific salt bridge. (C) εS280A transforms a key M2-M3 loop residue to the γ equivalent. (D) All mutants reduced AChR surface expression by ^125^–I-αBuTx radioligand binding assay. (E) Burst duration histograms of ε to γ mutants, longest time constant (ms) equals 5.58±0.34, 11.28±0.73, and 4.17±1.05 for adult WT, foetal WT, and εD163E+εD173F respectively, n≥4.

## Discussion

### Conformational changes between desensitised and resting states

Comparisons between our desensitised and resting state structures reveal new insights into the conformational changes between desensitised and resting states. We found that uncoupling of ACh is associated with asymmetrical movements in the two α M2-M3 loops away from δ or ε loop F, with distinct effects on the interactions between αS266 and δE189 or αS266 and εE184 (Figure 3B-C). The importance of the larger structural change at the αS266-δE189 interaction is supported by experimental mutation αS266A reducing AChR activity by 4-fold^51^ and the FCCMS εE184K variant’s effects on channel kinetics^14^. In contrast, an δE189A experimental mutation did not alter AChR function^51^. Therefore, we propose that the α-δ and α-ε interactions are non-equivalent, despite the sequence and structural homology of the δ and ε subunits. Additionally, our observation of another inter-subunit hydrogen bond between αS191 of loop C and δ/ε loop F, near the bound ACh, agrees with previous reports that this interaction stabilises the closed conformation of the orthosteric site^26,28,50,52^. Thus, loop F is an important component of signal transduction between the orthosteric sites and the coupling domain.

### Structural insights into CMS causing variants

Inappropriate treatment of CMS will lead to disease deterioration and can be life-threatening. Thus, correct identification of AChR variants as gain-of-function or loss-of-function is critical. Current gold standard for such identification is through functional assays such as single channel electrophysiology, which requires specialised expertise and equipment that is not accessible to most clinics. Whilst *in silico* prediction may provide a faster alternative, current technologies such as AlphaMissense are insufficient for clinical application in CMS due to an accuracy of 64% for predicting AChR missense variants^61,62^. Our cryo-EM structures and characterisation of eight CMS variants provide valuable information to help construct such a tool in the future. For example, by mapping the positions of known FCCMS and SCCMS variants, we found the variants are distributed differently across the AChR domains (Figures 4 and 5). FCCMS variants can be found throughout the AChR, whilst SCCMS variants concentrate in the TMD^7^. We reported five kinetic CMS variants on M3 that likely affect helical packing (Figure 5E). Yet our static structures alone are insufficient to explain the pathogenic mechanism of all the new variants we identified, most notably, why αI49T causes SCCMS and reduced channel conductance (Figure 5A-C). Nevertheless, our findings provide an initial understanding for the mutants’ physical effects. Building from these results and insights may enable the categorisation of future *de novo* CMS variants in AChR.

### Muscle-type AChR contains ion-coordinating ATCUN motifs

While unexpected, our structures provided a discovery of metal binding at the N-terminus of the receptor’s α subunits. We observed the α_δ_ ATCUN motif in an ion-bound conformation on structures obtained from copper grids (Figure 6B, Figure S10A-B) while in an open conformation on gold grids (Figure 6C). Intriguingly, the bound conformation of ATCUN has not been observed in the *Torpedo* or bovine AChR structures despite the sequence conservation, and the bovine AChR samples were also prepared on copper grids^24,28^. We propose the proximity of Fab35 binding to the ATCUN motifs may help to stabilise the α N-terminus to enable resolving this structural motif. Moreover, sequence differences between human and ortholog receptors adjacent to the ATCUN motifs may influence the site’s affinity for divalent cations. In the case of the *Torpedo* receptor, an isoleucine at the equivalent position as αV75 could disfavour the closed conformation of residue H3. In the case of the bovine receptor, a glutamine instead of εR23 at the α_δ_-ε interface cause the same effect. To further support our structural modelling, the bound AChR ATCUN motif also bears striking resemblance of other ion-coordinating ATCUN structures, such as that of a Gly-Gly-His modified tripeptide^63^, with backbone RMSD of 0.22 Å, and c-Src-SH3 domain^64^, with backbone RMSD of 0.31 Å (Figure S10C-D). Importantly, the functional significance of this newly identified metal binding site unclear, as it is distinct from known divalent cation binding sites of other pLGICs^65–67^.

### Assembly of muscle-type AChR subunits *in vitro*

Observations of the α_3_ε_2_ and α_2_βδ_2_ pentamers in our over-expression system provide insights into AChR folding, assembly, and trafficking *in vitro* as well as diagnosis of myasthenia gravis and CMS. These varied stoichiometries are of particular medical importance as cell-based assays are routinely used as diagnostic tests to detect anti-AChR antibodies in myasthenia gravis and for functional categorisation of novel CMS variants^10,68^. Therefore, inconsistencies between the native tissue and experimental model system could lead to erroneous clinical guidance.

The existence of multiple AChR stoichiometries in recombinant cells highlights the complexity of receptor assembly. Notably, whilst chaperones such as RIC-3 and NACHO are required for the overexpression of certain neuronal AChRs^69^, no chaperones have been definitively associated with the muscle-type AChR. Therefore, it is unclear whether the observed α_3_ε_2_ and α_2_βδ_2_ stoichiometries are due to the absence of a necessary chaperone in the HEK293 cells, or reflects assemblies present but not previously identified at the neuromuscular junction. However, our single-channel recordings demonstrated α-β-δ containing receptors appeared to function as an ACh-gated channel (Figure 7E). Furthermore, ACh-elicited currents were also recorded from α-β-δ containing receptors expressed in *Xenopus* oocytes^70^, and zebrafish knocked-out for ε or γ subunits still produced synaptic currents at the neuromuscular junction^71^. This evidence suggests the α_2_βδ_2_ receptor may play a physiological role *in vivo*.

Our observation of the α_3_ε_2_ pentamer suggests that AChR can form stoichiometries that have not been reported previously. The presence of these various stoichiometries has implications for the possible assembly mechanisms of this pLGIC^72^. However, it is not possible to infer all the *in cellulo* subunit assemblies in this dataset, as the necessary protein purification by the bromoacetylcholine affinity column biases the final sample to channels with at least one ACh binding site.

## Conclusion

Here, we demonstrated the recombinant overexpression of functional human muscle-type AChR using a stable cell line in quantities sufficient for structural biology. Having captured the resting and desensitised states of AChR by cryo-EM, our results reveal the receptor’s structural changes when converting from desensitised to resting state, in agreement with findings from previous ortholog structures^21–28^. Furthermore, our structures illuminate the underlying molecular pathophysiology of eight previously unstudied CMS variants. Finally, we identified a putative metal-binding ATCUN motif at the α subunit’s N-terminus, which may represent a possible regulatory site for AChR modulation by divalent cations. Ultimately, our models and results provide a new basis for understanding the structure-function relationship of the human AChR, and may help explain the molecular pathogenic mechanism of more CMS variants in the future.

## Methods

### Cell line generation and selection

Lentivirus was generated by first transfecting HEK293 cells with the AChR gene-containing TLCV2 plasmids, along with 2^nd^ or 3^rd^ generation lentiviral packaging and envelope plasmids (pHR_SIN, pCMV delta8.91/psPAX2, pMDG/pMD2.G). Cell culture medium consisted of Freestyle 293 medium (Gibco, 12338018) with 0.1% v/v penicillin-streptomycin-amphotericin B mix (PSA) (Lonza #17-745E). Lentivirus particles were harvested from the supernatant on day 4. To generate cell lines stably expressing AChR, on day 1 of cell line generation, 3 mL of Expi293F cells at 0.3×10^6^ cells/mL with 5 mg/mL hexamidine bromide (Merck #H9268) were seeded in a 24-well block (VWR #734-1217). Lentivirus for each AChR subunit were then each added to the cells at a final concentration of 0.15×10^6^-0.3×10^6^ transducing units (TU) per mL to achieve an approximate multiplicity of infection (MOI) of 1. The block was covered with porous adhesive film (VWR #391-1251) and grown in a shaking incubator (Infors HT Multitron) for 2 days at 37°C, 250 rpm, 75% humidity and 8% CO_2_, before exchanging to fresh medium on day 3. On day 4, induced cells were selected by adding 2.5 mg/mL puromycin. The medium was replaced every 2-5 days with an increasing amount of puromycin, until it reached 10 mg/mL. Selection was considered complete when the negative control of untransfected Expi293F cells undergoing the same puromycin selection were all dead (day 15).

Cells were then selected for AChR expression by first inducing expression with 0.1 μg/mL doxycycline (Merck #D3072) and then maintaining the cells at 0.3-3×10^6^/mL until GFP fluorescence appeared (day 17). The cells were then separated by FACS, using cell sorters (BD Aria FUSION or Sony MA900 FACS) fitted with a 100 mm nozzle and fluorescent filters (405-450/50, 561-610/20, 640-710/50 and 488-525/50) to select for quadruple positive cells (tagBFP^+^mIFP^+^mCherry^+^EGFP^+^) expressing all AChR subunits. Single cells were seeded into 100 μL medium with 10% v/v foetal bovine serum (FBS, PAA Laboratories or Gibco #10062-147) in 96-well round bottom plates (Corning #3799). Cells collected from FACS were cultured at 37°C, 75% humidity and 8% CO_2_ in a static incubator, replacing half of the supernatant with fresh medium every 1-3 days until colonies contained hundreds of cells before being dissociated with trypsin-EDTA (Gibco # 15400054) and transferred to 48-well plates (Corning #3548). Once confluent, the cells were transferred to 24-well plates (Corning #3526), followed by suspension-adaptation by growing in 24-well blocks and incubating with shaking at 250 rpm, passaging every 1-3 days to maintain a density of 0.3-3×10^6^ cells/mL. A final selection for AChR surface –expression was performed by radioactive ^125^I-αBuTx-binding assay, selecting the cell line with the highest receptor surface expression for large-scale protein production.

### Generation of Fab35

Mab35 is an IgG1 antibody against the AChR-α subunit produced by a rat immunised with *Electrophorus* AChR^38^. Mab35-expressing hybridoma cells (ATCC #TIB-175) were cultured in growth medium consisting of Dulbecco modified Eagle Medium (DMEM) (Gibco) with 10% v/v FBS and 1% v/v penicillin-streptomycin (Gibco #15140122) at 37°C and 8% CO_2._ Cells were sub-cultured every 2 to 3 days to maintain a density of 0.1-1×10^6^ cells/mL and expanded from a T25 flask (Corning #430639) to a T175 flask (Corning #431080). When cell reached complete confluency, the flask was filled with ∼450 mL growth medium and incubated until the medium turned orange at 2-3 weeks, with the semi-adherent cells resuspended every 3-5 days. Fab35 was then purified according to published protocol^73^.

### AChR expression, purification and nanodisc reconstitution

The AChR-expressing cell line was grown in Freestyle 293 medium (Invitrogen) with 0.1% v/v penicillin-streptomycin-amphotericin B mix in disposable roller bottles (Greiner Bio-One #680668 or Corning #431198) or Erlenmeyer flasks (Jet Biofil #JBE500 or Corning #431145) in a shaking incubator at 37°C, 100-170 rpm, 75% humidity and 8% CO_2_. Cells were maintained at 0.3-3×10^6^ cells/mL. Protein was expressed by first seeding one litre of cells (1×10^6^/mL) into each roller bottle, and then inducing expression the following day by adding 0.1 μg/mL doxycycline and 5 mM sterile-filtered sodium butyrate (Merck #8451440100). Cells were then grown at 30°C, 170 rpm for 3 days, and harvested by centrifugation at 1,500 g for 20 min at 4°C. Cell pellets were flash frozen in liquid nitrogen and stored at -80°C until use. ^74^

Membrane preparation was performed according to a protocol by Stevens *et al*^75^. Briefly, cell pellets from a 10 L of cell culture were thawed on ice, resuspended in 200 mL low salt buffer (50 mM HEPES-NaOH pH 7.5, 10 mM NaCl, 5% v/v glycerol, 5 mM MgCl_2_ and EDTA-free protease inhibitors (Roche #04693132001)) using a Dounce homogeniser (Merck), then lysed by 2 passes through a C3 or C5 Emulsiflex homogeniser (Avestin) at 10,000-15,000 psi. Lysate was brought to 400 mL with low salt buffer, then ultracentrifuged at 150,000 g for 45 min in a Ti45 rotor (Beckman Coulter). The membrane pellet was then washed by resuspending in low salt buffer with a Dounce homogeniser, then the volume adjusted to 400 mL and pelleting by ultracentrifugation again. Two additional washes were performed in high salt buffer (50 mM HEPES-NaOH pH 7.5, 1 M NaCl, 5% v/v glycerol, 5 mM MgCl_2_ and EDTA-free protease inhibitors) before the pellet was resuspended in 50 mL extraction buffer (50 mM HEPES-NaOH pH 7.5, 250 mM NaCl, 5% v/v glycerol), flash frozen in liquid nitrogen, and stored at -80°C until use.

To purify AChR, prepared membrane was thawed and resuspended in extraction buffer using a Dounce homogeniser. All steps of protein purification and cryo-EM sample preparation performed at 4°C or on ice. Protease inhibitor and 1% DDM/0.1% CHS (Anatrace #D310LA, Merck #C6512) were then added and the volume adjusted to 400 mL, and the sample was stirred for 1 h. Solubilised lysate was clarified by ultracentrifugation for 1 h at 100,000 g, then the supernatant was nutated with 5 mL pre-equilibrated bromoacetylcholine (BAC) affinity resins for 2 h. Bromoacetylcholine (BAC) affinity resin was generated by coupling bromoacetylcholine bromide (Santa Cruz #sc-252517) to Affi-Gel 10 (BioRad) according to a protocol by Bhushan and McNamee^74^. Flow-through was removed by centrifugation at 600 g for 5-7 min, with the resin transferred to a gravity column and washed with 5 mL extraction buffer (with 0.026% w/v DDM/CHS). Protein was eluted with elution buffer (50 mM HEPES-NaOH pH 7.5, 250 mM NaCl, 5% v/v glycerol, 100 mM carbamylcholine, 0.026% w/v DDM/CHS).

For detergent-solubilised AChR samples, EGFP-positive fractions detected by a ChemiDoc (BioRad) were pooled and incubated overnight with Fab35 and α-bungarotoxin (Invitrogen #B1601) 1:3:3 molar ratio. Next day, the sample was concentrated to 500 μL with 100 kDa concentrator (Sartorious), and then purified by Size-Exclusion Chromatography (SEC) using an ÄKTA Pure FPLC system (Cytiva) with a Sepharose 6 Increase 10/300 GL column (Cytiva) in gel filtration buffer (20 mM HEPES-NaOH pH 7.5, 150 mM NaCl, 0.013% w/v DDM/CHS). EGFP-positive fractions were concentrated to 1-2 mg/mL and used immediately for making cryo-EM grids.

For nanodisc reconstitution, EGFP-positive elution fractions were exchanged into extraction buffer with PD-10 columns (Cytiva) to remove detergent, then concentrated to 1 mL. Soy polar lipid (Avanti Polar Lipids #850457) was added to the protein at a 1:240 molar ratio of protein to lipid using a 50 mM stock in DDM/CHS and incubated with the protein. After 15 min, MSP2N2 (Merck #MSP12) was added at 1:6 ratio of AChR to MSP2N2 and nutated overnight. Next day, Biobeads (BioRad # 1523920) were added at the rate of 4 mg/h which was gradually increased to 20 mg/h, with the used Biobeads removed after every 3-5 additions, until the total mass of Biobeads exceeded 6 times the mass of detergent in the sample. During the final incubation with Biobeads, Fab35 and αBuTx were added to the protein at an estimated molar ratio of 3:3:1 and the mixture was rotated overnight. Next day, the supernatant was concentrated to 500 μL and separated by SEC in gel filtration buffer without detergent (20 mM HEPES-NaOH pH 7.5, 150 mM NaCl). GFP-positive fractions were pooled and concentrated to 1-2 mg/mL, spun at 17,000 g for > 30 min to remove aggregates, and the supernatant was collected to make cryo-EM grids.

Preparation of the desensitised sample followed the same protocol as the nanodisc-reconstituted sample with αBuTx, except αBuTx was replaced by 1 mM ACh or 100 μM ACh, 200 μM fluoxetine and 250 μM PAM^76^, which were incubated for > 1 h with the protein before sample vitrification.

### In-gel digest mass spectrometry

The presence of AChR subunits in Coomassie-stained SDS-PAGE of purified samples was confirmed by tryptic digestion and tandem mass spectrometry (LC-MS/MS) at the Centre for Medicines Discovery (CMD) Mass Spec Facility. Coomassie-stained SDS-PAGE gel bands were excised using a gel cutting tip (GeneCatcher, Web Scientific) and stored in 10% MeOH at 4°C. Prior to digestion, the methanol solution was removed and replaced with 100% acetonitrile for 2-5 min. The solution was then removed and replaced with 100 μL of 100 mM NH_4_HCO_3_ (pH 8.0). A further 1 μL of 1 M dithiothreitol was added and incubated at 56°C for 40 min, followed by the addition of 4 μL of 1 M iodoacetamide and the reaction incubated at ambient temperature in the dark for 20 min. Then, 1 μL of 1 M dithiothrietol, 200 μL of 100 mM NH_4_HCO_3_ and 1 μL of trypsin solution (sequencing grade, Sigma-Aldrich, 1 mg/mL in 0.01 M HCl) was added. Tryptic digestion proceeded at 37°C for 16 h and was terminated by the addition of 3 μL formic acid. Extracted peptides were analysed by TimsTOF 2 pro mass spectrometer (Bruker) with a nanoElute liquid chromatography system (Brucker), proteomics analysis was performed in MASCOT (Matrix Science).

### Cryo-EM sample preparation

The detergent-solubilised αBuTx-bound dataset was prepared on a UltrAuFoil R1.2/1.3 300 mesh gold grid, remaining three nanodisc-reconstituted datasets were prepared on Quantifoil R1.2/1.3 200 or 300 mesh copper grids. EM grids were glow-discharged for 60 s in air. Samples were vitrified using a Vitrobot Mark IV (FEI/Thermo Scientific) at 4°C and 100% humidity by applying to the grid 2.5-3 µL freshly prepared protein sample at 2-4 mg/mL, blotting for 2-4.5 s with a blot force of –10, and then immediate plunge freezing in liquid ethane.

### Cryo-EM data collection

Data collection parameters are summarised in Table S2.

For the αBuTx-bound detergent-solubilised dataset, a total of 18672 movies were recorded on a Titan Krios microscope at Oxford Particle Imaging Centre (OPIC, Oxford) using EPU control software (Thermo Scientific) and a Falcon 4i direct electron detector with a target defocus range of -1.2 to -2.4μm, a dose rate of 9.81 e^−^/pixel/s, and a total dose of 50 e^−^/Å^2^. Data were collected at a nominal magnification of 130,000x, corresponding to a calibrated pixel size of 0.932Å.

For the αBuTx-bound nanodisc dataset, a total of 30,556 movies were collected on a Titan Krios microscope at Central Oxford Structural Molecular Imaging Centre (COSMIC, Oxford) using EPU control software (Thermo Scientific) and a Gatan K3 detector with an energy filter slit width set to 20 eV. Data were recorded at a magnification of 105,000x, corresponding to a calibrated pixel size of 0.832 Å, using a target defocus range of -0.8 to -2.2 μm, a dose rate of 11.32 e^−^/pixel/s and a total dose of 46 e^−^/Å^2^. Movies were dose fractionated into 45 frames.

For the ACh-bound (100 μM) nanodisc dataset, 13,868 movies were recorded on Titan Krios with a Falcon 4i camera at Electron Bio-Imaging Centre (KriosIII, Diamond-eBIC, Harwell) using EPU control software (Thermo Scientific) at a nominal magnification of 130,000x for a calibrated pixel size of 0.921 Å. The grid was imaged using a target defocus range of -0.8 to - 2.4 μm, a dose rate of 6.64 e^−^/pixel/s, and total dose 50 e^−^/ Å^2^. The Selectris X energy filter slit width was set at 10 eV.

For the ACh-bound (1 mM) ‘desensitised’ nanodisc dataset, a total of 19,925 movies were collected on a Titan Krios microscope at Central Oxford Structural Molecular Imaging Centre (COSMIC, Oxford) using EPU control software (Thermo Scientific) with a Gatan K3 detector using a nominal magnification of 130000x for acalibrated pixel size 0.65 Å. The energy filter slit width was set to 20 eV, the target defocus range was -1.0 to -2.2 μm, with a dose rate of 13.90 e^−^/pixel/s, and a total dose of 52.2 e^−^/Å^2^. Movies were dose fractionated into 50 frames.

### Cryo-EM image processing

Data processing workflows are summarised in Figures S2, S5, S7-S8. Data were processed with cryoSPARC (v3.3.1 / 4.4.1)^77^ and RELION (v3.1 / v5.0)^58^. Electron Event Representation (EER) movies were fractionated into frames with a dose of 1 e^−^/Å^2^ for the detergent solubilised image set and 0.476 e^−^/Å^2^ for the 100 μM ACh image set, which were collected on a Falcon 4i detector, prior to motion-correction. Motion correction was carried out using cryoSPARC’s patch motion correction for the detergent-solubilised sample, SIMPLE for the αBuTx-bound AChR in nanodisc, and RELION for the two remaining ACh-bound datasets; this was followed by patch CTF-correction, particle-picking and 2D classification in cryoSPARC^58,77–79^. Particles from well resolved 2D classes were used for *ab initio* model generation followed by rounds of heterogeneous refinement to isolate particles with either one or two bound Fab35 molecules. Initial processing of all datasets highlighted the presence of several non-α_2_βδε stoichiometries so further downstream processing focussed on the 2Fab class. After non-uniform refinement, aligned particles were exported from cryoSPARC to RELION using *csparc2star*.*py* from the PYEM package^80^. Particles from the nanodisc-reconstituted datasets underwent per particle CTF refinement and Bayesian polishing in RELION. All particle sets also underwent static 3D classification in RELION using focussed masks generated from Alphafold2^81^ initial models using UCSF ChimeraX^82^ (Table S2, Figures S2,S5, S7-S8). Less rounds of static 3D classification were performed on the nanodisc image sets due to their overall preferential particle orientational bias. After static 3D classification, final particle sets were reimported into cryoSPARC for a final round of non-uniform refinement to yield the final reconstructions^83^. Subunit stoichiometries were checked at each stage of data processing using unsupervised automated model building with ModelAngelo^57^, with the longest traced sequence of each subunit identified using the ‘Blast Protein’ tool within UCSF ChimeraX^81,84^.

### Model building and refinement

Coordinates for αBuTx and Fab35 were derived from PDB entry 5HBT^73^. Coordinates for AChR subunits were derived from the AlphaFold2 model database, which was used to generate the initial models^81^. Initial models were fitted into the maps using UCSF ChimeraX^82^ and then adjusted and real-space refined in COOT^85^. In addition, models built using ModelAngelo^57^ were used to guide manual model building in COOT. Residue numbering for AChR subunits start after the signal peptide sequences, in accordance with the established AChR nomenclature where αL251 is situated at the 9’ midpoint position of the pore^23^. Rebuilt models were then refined against appropriately *B*-factor sharpened maps in ISOLDE^86^ before final global real-space refinement in PHENIX 1.21^87^ using appropriate reference model and geometric restraints. The αBuTx peptides were modeled into the low occupancy density of the detergent-solubilised sample by docking αBuTx from 5HBT into a blurred map, and their location confirmed in the better resolved αBuTx density of the nanodisc dataset. For the ACh-bound structures, an initial model was built using the map from 100 µM ACh data as this was visually superior to the 1 mM ACh map. ACh was well-defined in the Coulomb potential map and could be placed unambiguously. Ligand restraints were generated using GRADE2^88^ and the modelled ACh conformation was validated with MOGUL^89^. In all structures, a disulphide bond was apparent and modelled between αC192 and αC193 in loop C although there was some evidence of partial S-S reduction particularly in the resting nanodisc structure. In general, the M1-M4 transmembrane helices were worse resolved in all the nanodisc datasets (Figure S6) and it was not possible to unambiguously assign the position of all sidechains in the TMD, thus, the ambiguous sidechains were built to polyalanine. N-glycan structures were validated with Privateer assuming hybrid chains and human expression system^90,91^. Pore profiles were generated from structures using the HOLE^92^ implementation in COOT^85^ after reinstating any truncated side chains using ICM Browser Pro (Molsoft L.L.C).

To highlight significant non-protein density, mask-normalised F_o_-F_c_ difference maps were generated by refining the final model with all non-protein atoms removed using Refmac Servalcat^93^ within the CCPEM-1.6.0 suite and the refinement mask from cryoSPARC’s non-uniform refinement to normalise the sigma levels. Prior to difference map calculation, models were refined for 30 cycles against sharpened half maps using jelly-body restraints.

Sequence-independent structural superpositions of the relevant AChR chains were carried out in pyMOL (The PyMOL Molecular Graphics System, Version 3.0 Schrödinger, LLC) using the ‘super’ command, and the output RMSD values are for all equivalent atoms after structural outlier rejection.

### CMS patient variants

Suspected cases of congenital myasthenic syndrome are referred to the ‘Oxford highly specialised clinical service for CMS’ for genetic analysis and/or AChR functional assessment to determine pathogenicity. Variants in the AChR subunit genes identified by next-generation whole genome/exome sequencing, and confirmed in the patient DNA by Sanger sequencing, were introduced into respective AChR subunit cDNAs by site-directed mutagenesis. Following mutagenesis, constructs were revalidated by Sanger sequencing and then used in functional or electrophysiological studies (see construct design, electrophysiology and radioactive ^125^I-αBuTx-binding assay).

### Construct design

Complementary DNAs (cDNA) encoding human wildtype AChR α1-(P3A negative isoform), β1-, and δ-subunits in pcDNA3.1-hygro were generated previously^94^. The ε_EGFP_ construct was generated by inserting EGFP from pEGFP-N1 (BD Biosciences) into the M3-M4 loop of wildtype ε sequence using the *Sfi*I restriction site^36^. The doxycycline-inducible lentiviral TLCV2 vector (AddGene: 87360) was first modified by a) removing the Cas9 sequence from the original vector, b) inserting a KOZAK sequence, and b) inserting an internal ribosome entry site (IRES2) followed by fluorescent protein cDNA of tagBFP, mCherry, mIFP, or EGFP ^95–98^, before the AChR cDNA was cloned into the vector between the KOZAK and IRES2 to create the co-expression constructs: α-tagBFP, β-mIFP, and δ-mCherry. The construct for the ε_EGFP_ fusion protein contained no fluorescent protein following IRES2. Final constructs were verified by Sanger sequencing.

### Electrophysiology

Recordings were made from HEK293 cells 48 h after transfection with AChR subunit cDNAs in pcDNA3.1 vector, withEGFP-N1 (Clonetech) included as a marker of transfection. Recordings were performed in the cell-attached patch configuration at 20-22°C^99^. The cells were bathed in a solution containing 150 mM NaCl, 2.8 mM KCl, 2 mM MgCl_2_, 1 mM CaCl_2_, 10 mM HEPES-NaOH, 10 mM glucose, at pH 7.4. The pipette solution was the same as bath solution, except glucose was omitted and ACh added to 100 nM or 500 nM (for δD180N variant only). Single-channel currents were amplified with an Axopatch 200B amplifier (Molecular Devices, Sunnyvale, California), and sampled at 100 kHz, initially filtered at 5 kHz (−3 dB, Bessel filter), with a resolution of 50 μs. Burst duration recordings were made with the pipette potential set at +80 mV. For the determination of channel conductance, pipette holding potentials was varied from +20 mV to +120 mV. Channel transitions were detected by 50% amplitude threshold crossings (pClamp10). Bursts were defined as groups of openings separated by closed intervals longer than a critical duration (t_crit_). t_crit_ was determined for each patch using Equation (1)^100^:

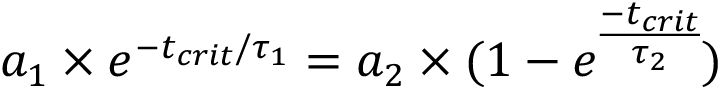

where a and τ indicate area and time constant for within and between bursts. Histograms of burst duration were fitted to the sum of exponentials by maximum log likelihood. Wildtype α_2_βδε channels and most mutants were typically best fit by the sum of 3 exponential functions, though aI49T that was best fit by 4 exponentials. In some cases, insufficient channel activity was recorded from individual patches; in those cases, recorded burst from multiple patches were combined and then fit by the sum of exponentials. In these cases, the fitting error is reported. The τ values of the population with longest duration are reported and compared with WT values.

### Radioactive ^125^I-αBuTx-binding assay for measuring AChR surface expression

HEK293 cells were grown in DMEM (Gibco) supplemented with 10% FCS and 1% PSA at 37°C and 5% CO_2_. Cells were seeded at 3×10^5^ cells per well of a 6-well plate and transfected the following day with 3 µg DNA per well with a ratio of 2:1:1:1 of human muscle AChR α1, β1, δ, and ε subunits. Forty eight hours post-transfection, ^125^I-αBuTx (Revvity/PerkinElmer) was added to the cells at 1×10^6^ counts per minute (cpm) per well in 1 mL phosphate buffered saline (PBS, Oxoid #BR0014G) with 1 mg/mL bovine serum albumin (Sigma-Aldrich) and incubated at room temp for 1 h. Each well of adherent cells was washed three times for 5 min with PBS, removed from the plate using 0.5 mL lysis buffer (10 mM Tris-HCl, 100 mM NaCl, 1 mM EDTA, 1% Triton X-100, pH 7.5), and levels of gamma radiation (in cpm) were recorded by a Cobra-II Auto Gamma Counter (Packard).

To analyse the εS235A CMS variant, an extra immunoprecipitation step was performed on the transfected cells to eliminate the contaminating effects of αβδ receptors. Cells were labelled with ^125^I-αBuTx, washed in PBS, and removed from plates in lysis buffer as described before. This ensures only receptors on the cell surface were labelled. For immunoprecipitation, the lysate was incubated with 5 µL of the ε subunit-specific polyclonal rabbit antiserum^101^ overnight at 4°C. Following day, 75 µL of goat anti-rabbit IgG (Phoenix Pharmaceuticals, #RK-GAR) was added for 2 h at room temperature with rotation.

Precipitate was spun down at 15,000-20,000 g for 5 min and analysed for the residual gamma radiation.

## Acknowledgements

We thank neuromuscular clinicians for referring AChR sequence variants to the ‘Oxford highly specialised clinical service for CMS’ for functional analysis. We thank Hayley Spearman for conducting initial surface expression experiments for the βV296A mutant; Professor Angela Vincent and Professor Alex Bullock as well as members of the Yin Dong and David Sauer labs for useful discussions; Dr Karin Rödström for training A.L. to purify proteins, Dr Philip Hublitz and his team at the Genome Engineering Facility in the WIMM for cloning the AChR lentiviral constructs; Dr Ryan Beveridge at the Virus Screening Facility in the WIMM for generating the lentiviral particles and calculating the virus titre; Craig Waugh, Dr Paul Sopp and Kevin Clark for assisting with cell sorting at the WIMM FACS facility; Dr Rod Chalk and Dr Kavya Clement for assisting with LC-MS/MS analysis; Professor Brian Marsden, Elizabeth Maclean (Centre for Medicines Discovery) for cryo-EM support. Cryo-EM datasets were collected at the Oxford Particle Imaging Centre (OPIC), Central Oxford Structural Molecular Imaging Centre (COSMIC), and the electron Bio-Imaging Centre (eBIC) inside Diamond Light Source (BAG proposal bi28713). We acknowledge funding from the Wellcome Trust (102161/Z/13/Z, studentship to A.L.), UK Medical Research Council (MRC) (MR/S007180/1 and MR/Z504099/1 to Y.Y.D.), and MRC MR/Y012623/1 to A.L., Y.Y.D. and D.B.S. D.B.S. and A.C.W.P. were supported by the Innovative Medicines Initiative 2 Joint Undertaking (JU) under grant agreement No 875510 (EUbOPEN). The JU receives support from the European Union’s Horizon 2020 research and innovation program and EFPIA and Ontario Institute for Cancer Research, Royal Institution for the Advancement of Learning McGill University, Kungliga Tekniska Högskolan, Diamond Light Source Limited. OPIC is an Instruct-ERIC centre funded by Wellcome Trust JIF award (060208/Z/00/Z) and equipment grant (093305/Z/10/Z). COSMIC is supported by the Wellcome Trust (grant no. 201536), the EPA Cephalosporin Trust and a Royal Society/Wolfson Foundation Laboratory Refurbishment grant (no. WL160052). Molecular graphics and analyses performed with UCSF ChimeraX, developed by the Resource for Biocomputing, Visualization, and Informatics at the University of California, San Francisco, with support from National Institutes of Health R01-GM129325 and the Office of Cyber Infrastructure and Computational Biology, National Institute of Allergy and Infectious Diseases.

## Lead Contact

Further information and requests for resources and reagents should be directed to and will be fulfilled by the Lead Contact, Yin Yao Dong.

## Materials Availability

All unique reagents generated in this study are available from the Lead Contact with a completed Materials Transfer Agreement.

## Data and code availability

The cryo-EM maps were deposited to the EM Data Bank under accession codes EMD-51568 (αBuTx-detergent), EMD-51569 (αBuTx-nanodisc), EMD-51570 (100 μM ACh-nanodisc), and EMD-51571 (1 mM ACh-nanodisc). The atomic models were deposited to the PDB under accession codes 9GU0 (αBuTx-detergent), 9GU1 (αBuTx-nanodisc), 9GU2 (100 μM ACh-nanodisc), and 9GU3 (1 mM ACh-nanodisc).

## Author contributions

Y.Y.D. and A.L. designed the experiments. A.L. performed the cell line generation, protein production and cryo-EM sample preparation, and drafted the manuscript. A.L., A.C.W.P. and G.C. conducted cryo-EM screening and data collection. A.L. and A.C.W.P. performed cryo-EM data analysis, model building, and structural analysis. R.W. performed single channel electrophysiology. S.M. and W.L. carried out mutagenesis cloning and surface expression assay. J.P. led the ‘Oxford Congenital MS’. Y.Y.D., D.B.S., and D.B. obtained funding and supervised the research. A.L., D.B.S., A.C.W.P. and Y.Y.D. revised the manuscript with input from all authors.

