## Supplementary information for "Structures of the human adult muscle-type nicotinic receptor in resting and desensitised states"

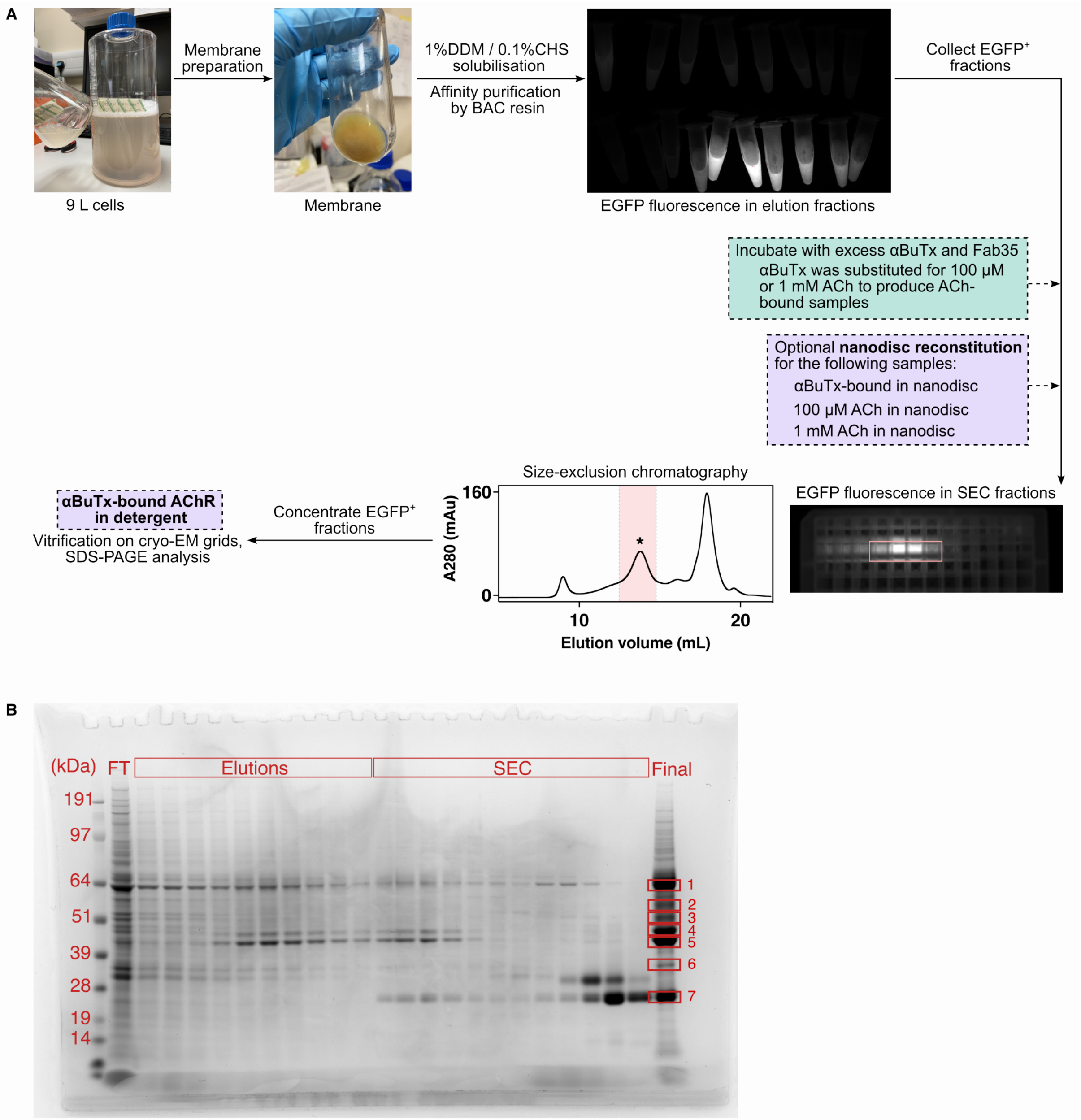


**Figure S1**: Expression and purification of AChR from stable cell line. (A) Purification workflow of αBuTx-bound AChR in detergent. Lilac, optional nanodisc reconstitution step was performed for the following datasets: αBuTx-bound in nanodisc, 100 μM ACh in nanodisc and 1 mM ACh in nanodisc. Turquoise, purified AChR was incubated with excess ACh instead of αBuTx to produce the ACh-bound samples. (B) SDS-PAGE of purified AChR with added Fab35, bands 1-7 (in boxes) were excised for protein identification (see Table S1).

**Table S1: Protein identification by in-gel digest mass spectroscopy**

| Band | Protein Score | Coverage (%) | emPAI | #Significant Sequence | Average Mass | Name | Description |
|---|---|---|---|---|---|---|---|
| 1 | 10356 | 41 | 1.68 | 20 | 54662 | ACHE_HUMAN | Acetylcholine receptor subunit epsilon OS=Homo sapiens OX=9606 GN=CHRNE PE=1 SV=2 |
| 2 | 4214 | 46 | 1.5 | 20 | 58858 | ACHD_HUMAN | Acetylcholine receptor subunit delta OS=Homo sapiens OX=9606 GN=CHRND PE=1 SV=1 |
| 3 | 3859 | 37 | 1.38 | 19 | 58858 | ACHD_HUMAN | Acetylcholine receptor subunit delta OS=Homo sapiens OX=9606 GN=CHRND PE=1 SV=1 |
| 4 | 7597 | 37 | 1.24 | 17 | 56663 | ACHB_HUMAN | Acetylcholine receptor subunit beta OS=Homo sapiens OX=9606 GN=CHRNB1 PE=1 SV=3 |
| 5 | 13193 | 44 | 1.68 | 19 | 51805 | ACHA_HUMAN | Acetylcholine receptor subunit alpha OS=Homo sapiens OX=9606 GN=CHRNA1 PE=1 SV=3 |
| 6 | 1657 | 36 | 1.07 | 14 | 51805 | ACHA_HUMAN | Acetylcholine receptor subunit alpha OS=Homo sapiens OX=9606 GN=CHRNA1 PE=1 SV=3 |
| 7 | 1473 | 28 | 1.41 | 4 | 11594 | KACB_RAT | Ig kappa chain C region, B allele OS=Rattus norvegicus OX=10116 PE=1 SV=1 |

### Table S2: Cryo-EM data collection, processing, refinement and validation statistics

| | **αBuTx-detergent** (EMD-51568) (PDB 9GU0) | **αBuTx-nanodisc** (EMD-51569) (PDB 9GU1) | **100 µM ACh-nanodisc** (EMD-51570) (PDB 9GU2) | **1 mM ACh-nanodisc** (EMD-51571) (PDB 9GU3) |
|---|---|---|---|---|
| **Data collection and processing** | | | | |
| Magnification | 130,000 | 105,000 | 130,000 | 130,000 |
| Voltage (kV) | 300 | 300 | 300 | 300 |
| Electron exposure (e–/Å$^2$) | 50 | 46 | 50 | 52.2 |
| Defocus range (µm) | -1.2 to -2.4 | -0.8 to -2.2 | -0.8 to -2.4 | -1.0 to -2.2 |
| Pixel size (Å) | 0.932 | 0.832 | 0.921 | 0.65 |
| Symmetry imposed | C1 | C1 | C1 | C1 |
| Initial particle images (no.) | 696904 | 795093 | 933302 | 1713360 |
| Final particle images (no.) | 31022 | 319381 | 175800 | 434976 |
| Map resolution (Å) | 2.96 | 2.48 | 2.73 | 2.64 |
| FSC threshold | 0.143 | 0.143 | 0.143 | 0.143 |
| Map resolution range (Å) | 2.5 – 6.5 | 2.3 – 6.0 | 2.5 – 6.0 | 2.6 – 6.0 |
| **Refinement** | | | | |
| Initial model used (PDB code) | Alphafold2 | 9GU0 | Alphafold2/ModelAngelo | 9GU2 |
| Model resolution (Å) | 3.31 | 2.74 | 2.88 | 2.91 |
| FSC threshold | 0.5 | 0.5 | 0.5 | 0.5 |
| Map sharpening *B* factor (Å$^2$) | -20 | -25 | -40 | -25 |
| Model composition | | | | |
| Non-hydrogen atoms | 22876 | 21896 | 21623 | 19563 |
| Protein residues | 2956 | 2740 | 2748 | 2564 |
| Ligands | 0 | 0 | 20 | 20 |
| *B* factors (Å$^2$) | | | | |
| Protein | 113.08 | 128.10 | 114.50 | 162.27 |
| Ligand | - | - | 48.01 | 74.35 |
| R.m.s. deviations | | | | |
| Bond lengths (Å) | 0.002 | 0.005 | 0.002 | 0.003 |
| Bond angles (°) | 0.497 | 0.616 | 0.488 | 0.504 |
| Validation | | | | |
| MolProbity score | 1.23 | 1.32 | 1.34 | 1.32 |
| Clashscore | 2.46 | 3.63 | 3.38 | 4.47 |
| Poor rotamers (%) | 0.21 | 0.21 | 0.44 | 0.46 |
| Ramachandran plot | | | | |
| Favoured (%) | 96.76 | 97.08 | 96.65 | 97.55 |
| Allowed (%) | 3.24 | 2.92 | 3.35 | 2.45 |
| Disallowed (%) | 0 | 0 | 0 | 0 |
| Overall Q-score | 0.4970 | 0.5060 | 0.4970 | 0.4480 |

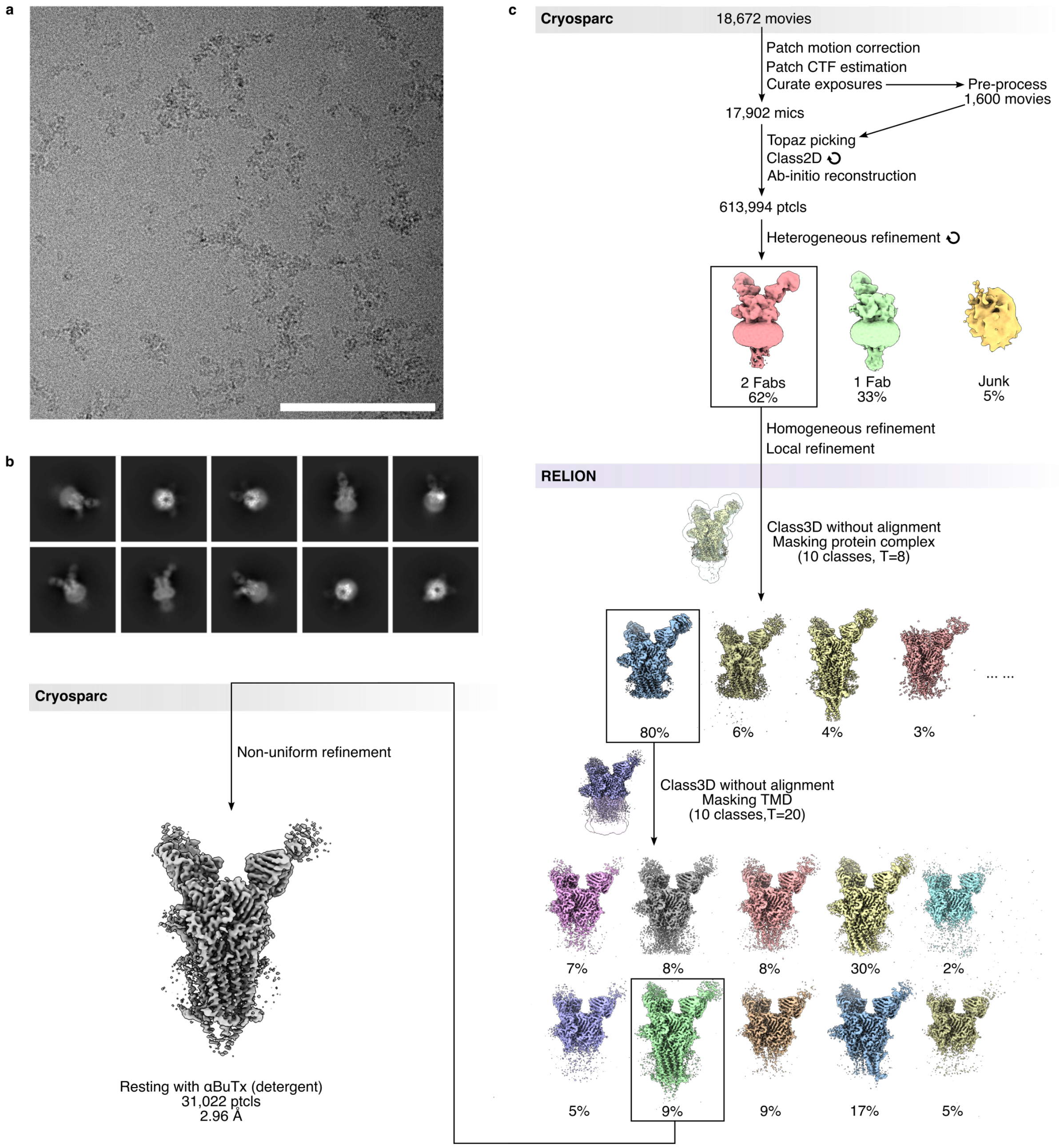


**Figure S2:** Cryo-EM data processing flowchart for resting state structure in detergent. (A) Representative micrograph, scale bar is 100 nm. (B) Representative 2D classes. (C) Data processing workflow. AChR particles were Topaz-picked from motion-corrected movies and subjected to 3D reconstruction and heterogeneous refinement in CryoSPARC v3.3.1[1-3]. The 2Fab particles were imported into RELION 3.1.3[4] for static Class3D with a mask covering the entire protein complex for the first round, and another mask covering the TMD for the second round. The particles were re-imported into CryoSPARC[5] for non-uniform refinement to produce a reconstruction map of 2.96 Å with 31,022 particles.

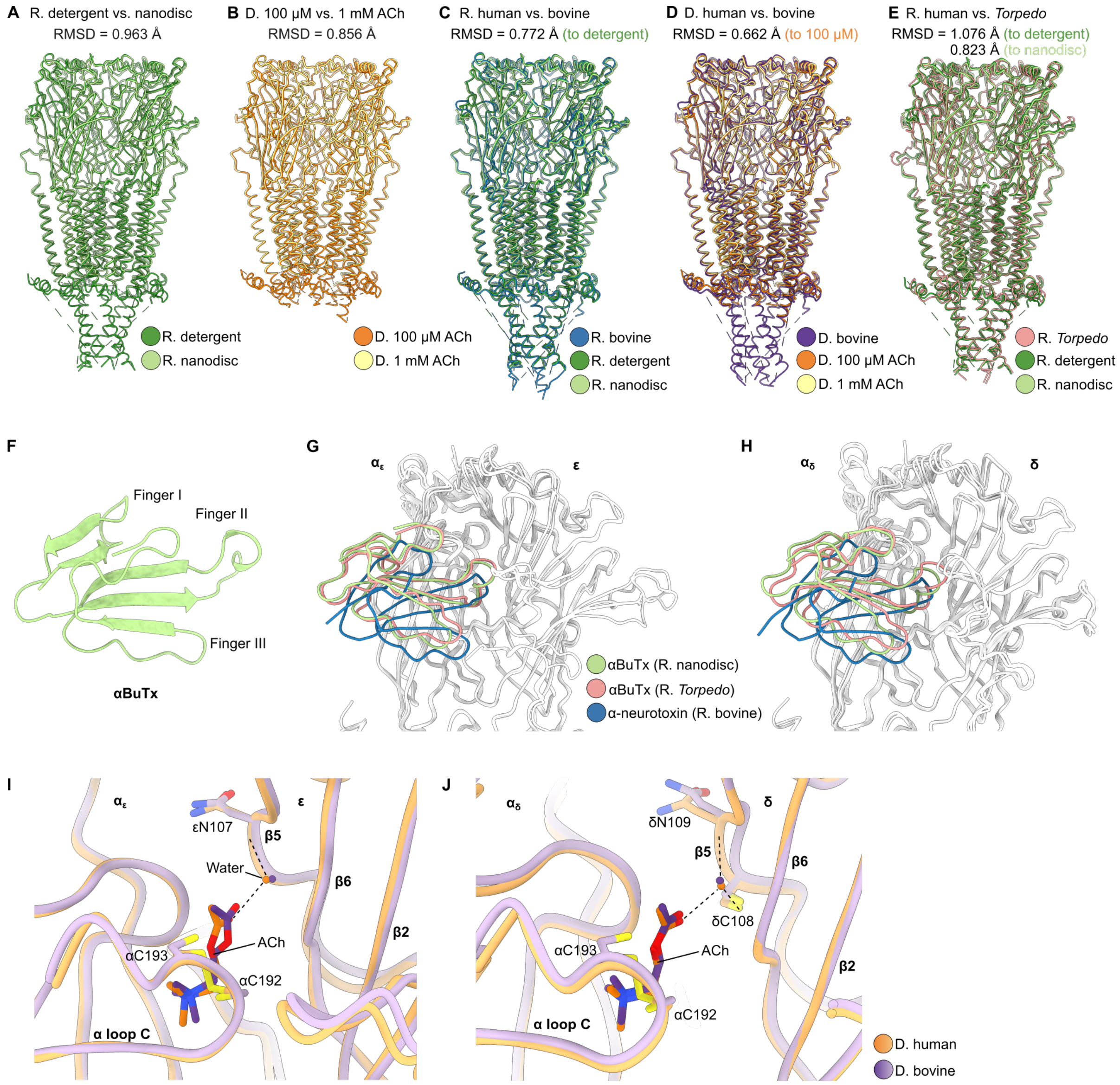


**Figure S3: Comparisons of αBuTx and ACh binding to ortholog structures**. (A-E) Structural overlay of two human resting state (R.) structures, two human desensitised (D.) structures, human versus bovine in the resting state (PDB: 9AVV), human versus bovine in the desensitised state (PDB: 9AWJ), and human versus *Torpedo* in the resting state (PDB: 6UWZ) [6,7]. All resting state structures are bound to α-neurotoxins. RMSD values (in Å) are 0.963, 0.856, 0.772, 0.662 and 1.076 (to detergent) or 0.823 (to nanodisc). (F) The αBuTx is a three-finger α-neurotoxin. (G-H) An engineered α-neurotoxin binds α-ε and α-δ interfaces in a different manner to αBuTx. (I-J) ACh adopts a different pose compared a bovine AChR structure (9AWJ) possibly due to different oxidation states of the αC192-αC193 disulphide bond.

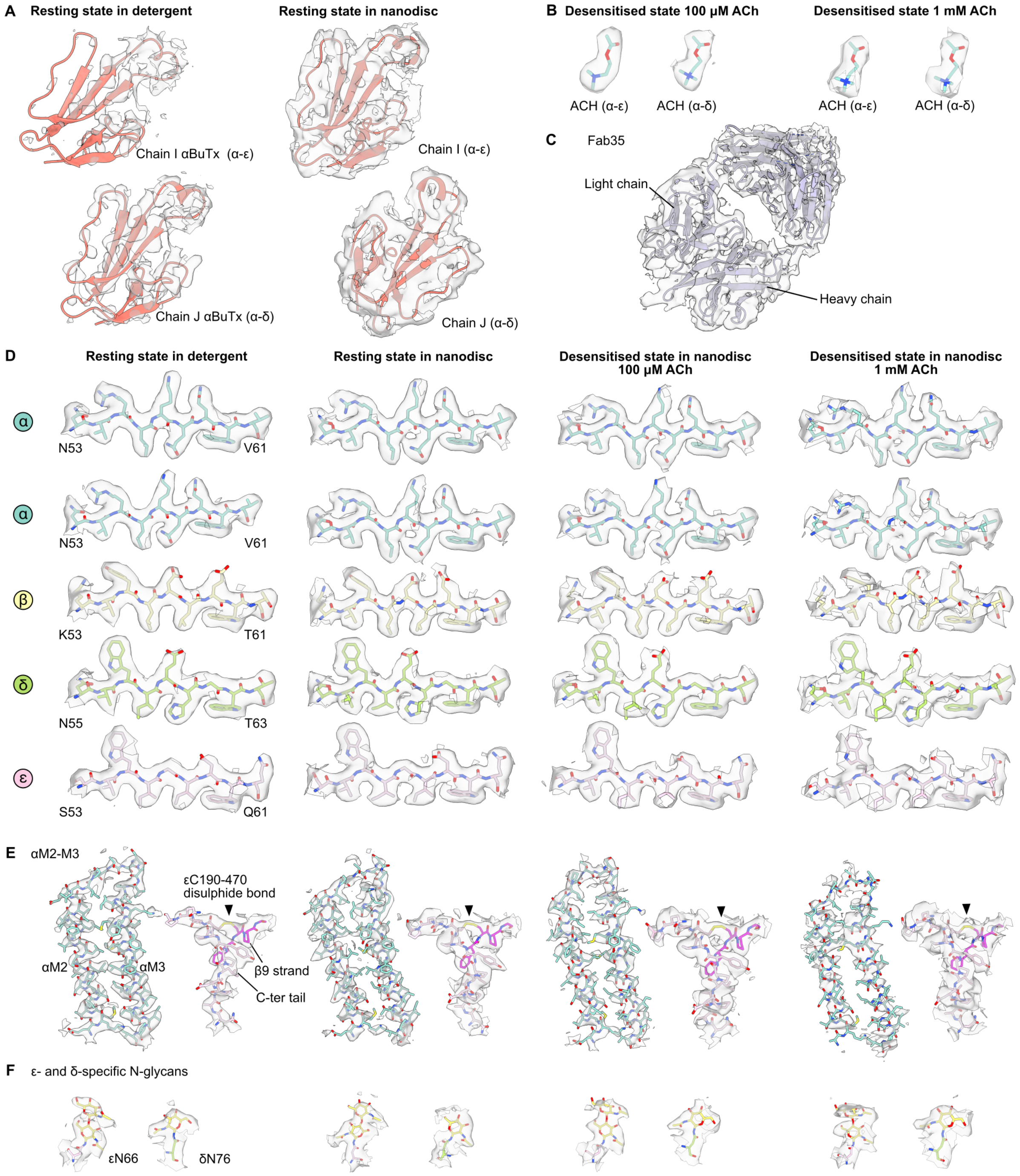


**Figure S4: Cryo-EM map quality.** Representative views of subunit signature sequence in ECD-β2 strand of the $\alpha_\varepsilon$ subunit (A), M2-M3 helices with ε C-terminal disulphide bond (εC190-εC470) (B), N-glycans (C), αBuTx (D), ACh (E) and Fab35 (F) are shown.

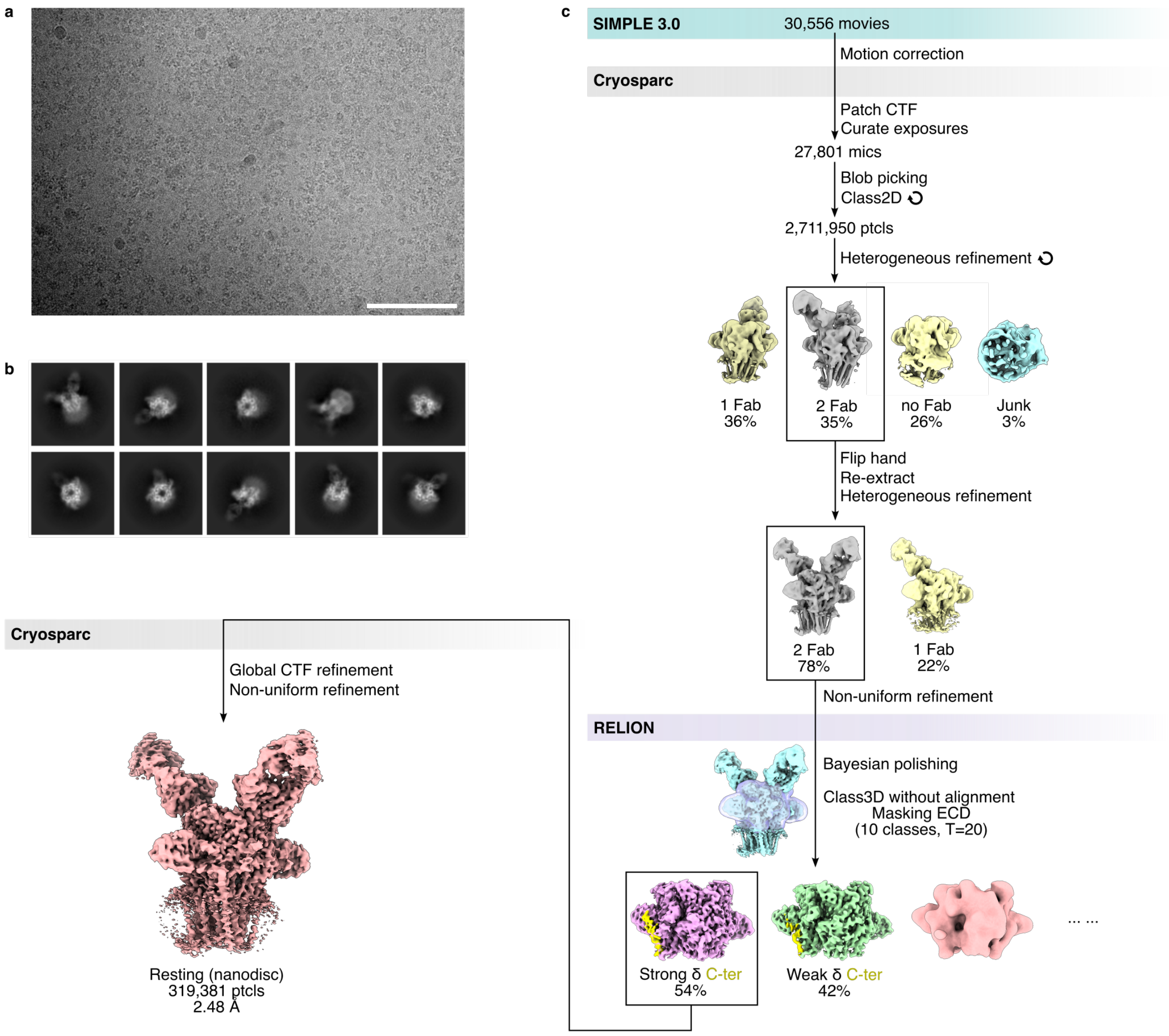


**Figure S5**: Cryo-EM data processing flowchart for resting state structure in nanodisc. (A) Representative micrograph, scale bar is 100 nm. (B) Representative 2D classes. (C) Data processing workflow. The classification steps were simplified compared to Figure S2 due to severe orientation bias. A mask covering the ECD was generated from initial model (see Methods) and used for RELION Class3D of 2Fab particles. This step selected for high-resolution particles with full αBuTx occupancy at both binding sites, as well as correctly aligned α-β-δ-α-ε subunits. Two high-resolution particle subclasses emerged, one with strong densities for the long δ C-terminal tail (54%) and the other with weak density for the δ signature motif (42%). The subclass with strong δ C-terminal tail density showing unambiguous subunit stoichiometry was selected for refinement, which produced a final reconstruction at 2.48 Å with 319,381 particles.

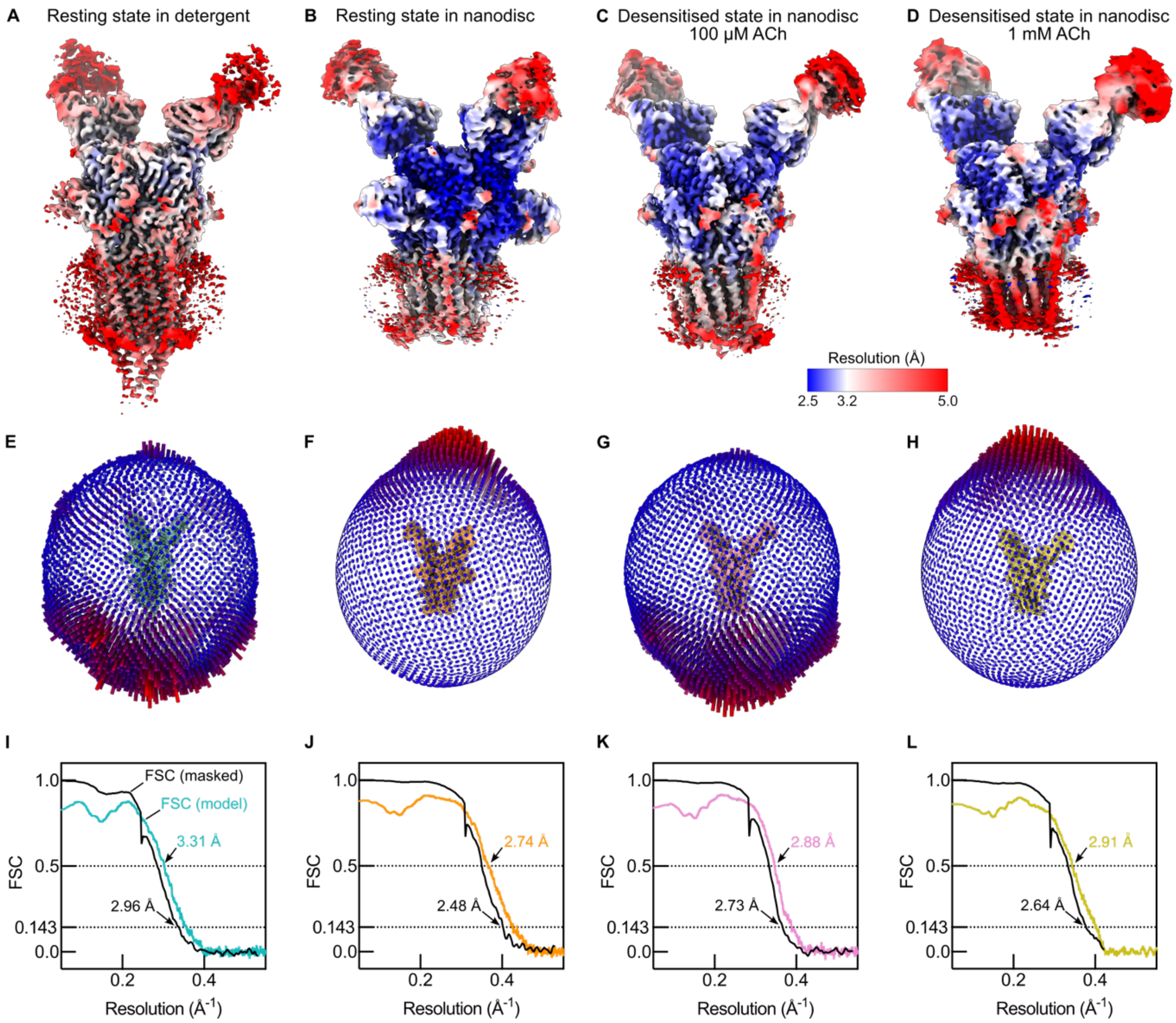


**Figure S6: Cryo-EM local resolution maps, angular distribution and FSC curves.** (A-D) Local resolution of cryo-EM maps. (E-H) Angular distribution of particles for each dataset. (I-L) Masked FSC curves (black) and map-to-model FSC curves (coloured).

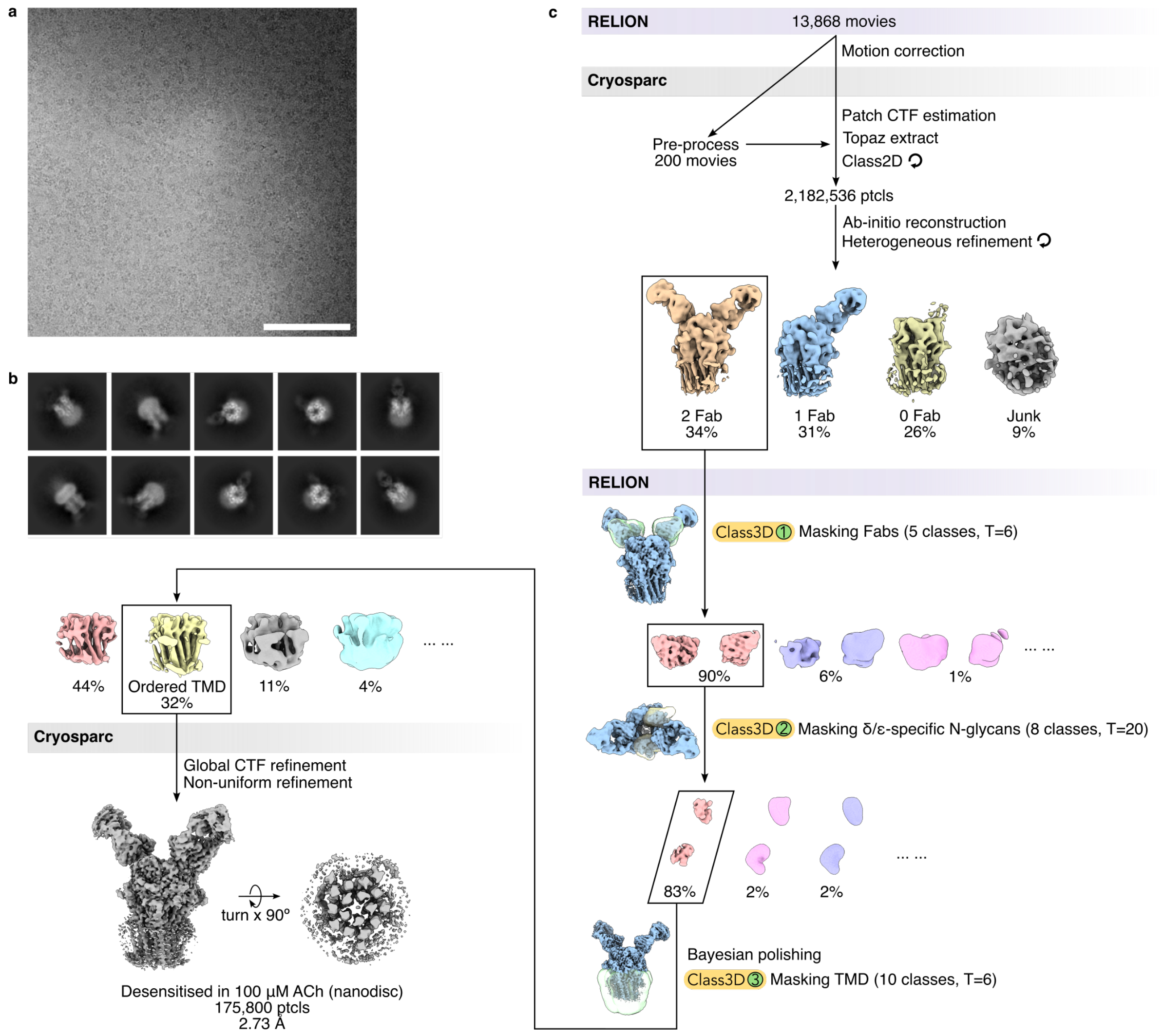


**Figure S7:** Cryo-EM data processing flowchart for desensitised state structure in nanodisc with 100 μM acetylcholine. (A) Representative micrograph, scale bar is 100 nm. (B) Representative 2D classes. (C) Data processing workflow. Two-Fab particles were initially isolated in CryoSPARC, then subjected to three rounds of RELION static Class3D. The first round involved a mask covering the two Fabs to select for particles with two α subunits, the second round used a different mask covering the δ/ε-specific N-glycans to ensure the resultant pentamer contains a δ or ε subunit at the correct position, and finally, the third round involved yet another mask covering the TMD to select for particle subclasses with ordered helices at the pore. From the final round of static class3D, a high-resolution particle subclass with ordered TMD (32%) emerged, and after refinement, it produced a 2.73 Å final reconstruction with 175,800 particles.

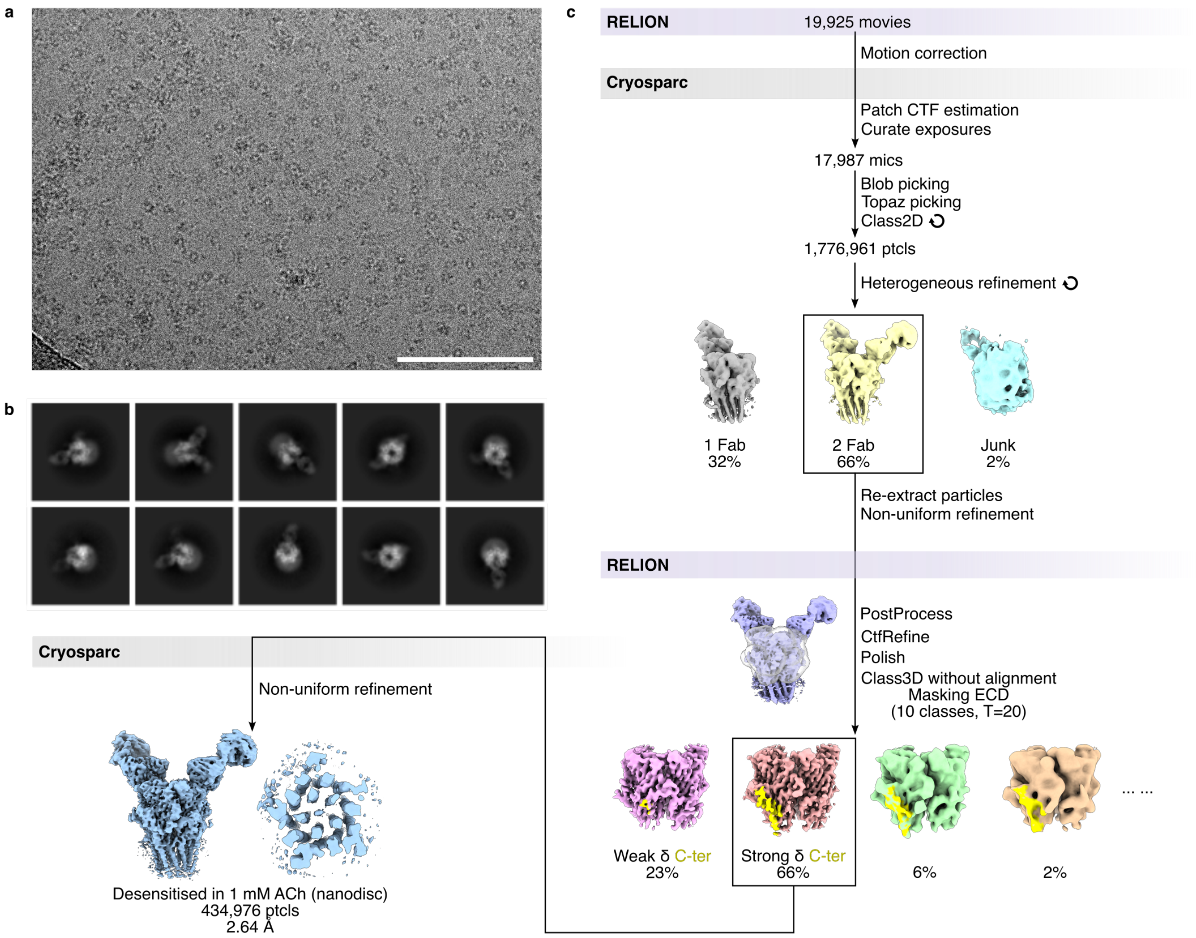


**Figure S8**: Cryo-EM data processing flowchart for desensitised state structure in nanodisc with 1 mM acetylcholine. (A) Representative micrograph, scale bar is 100 nm. (B) Representative 2D classes. (C) Data processing workflow. The classification steps were similar to Figure S5 due to severe orientation bias. A mask covering the ECD was used for static Class3D of 2Fab particles, which isolated a high-resolution particle subclass with strong densities for the long δ C-terminal tail (66%). Refinement of this particle subclass produced a final reconstruction at 2.64 Å with 434,976 particles.

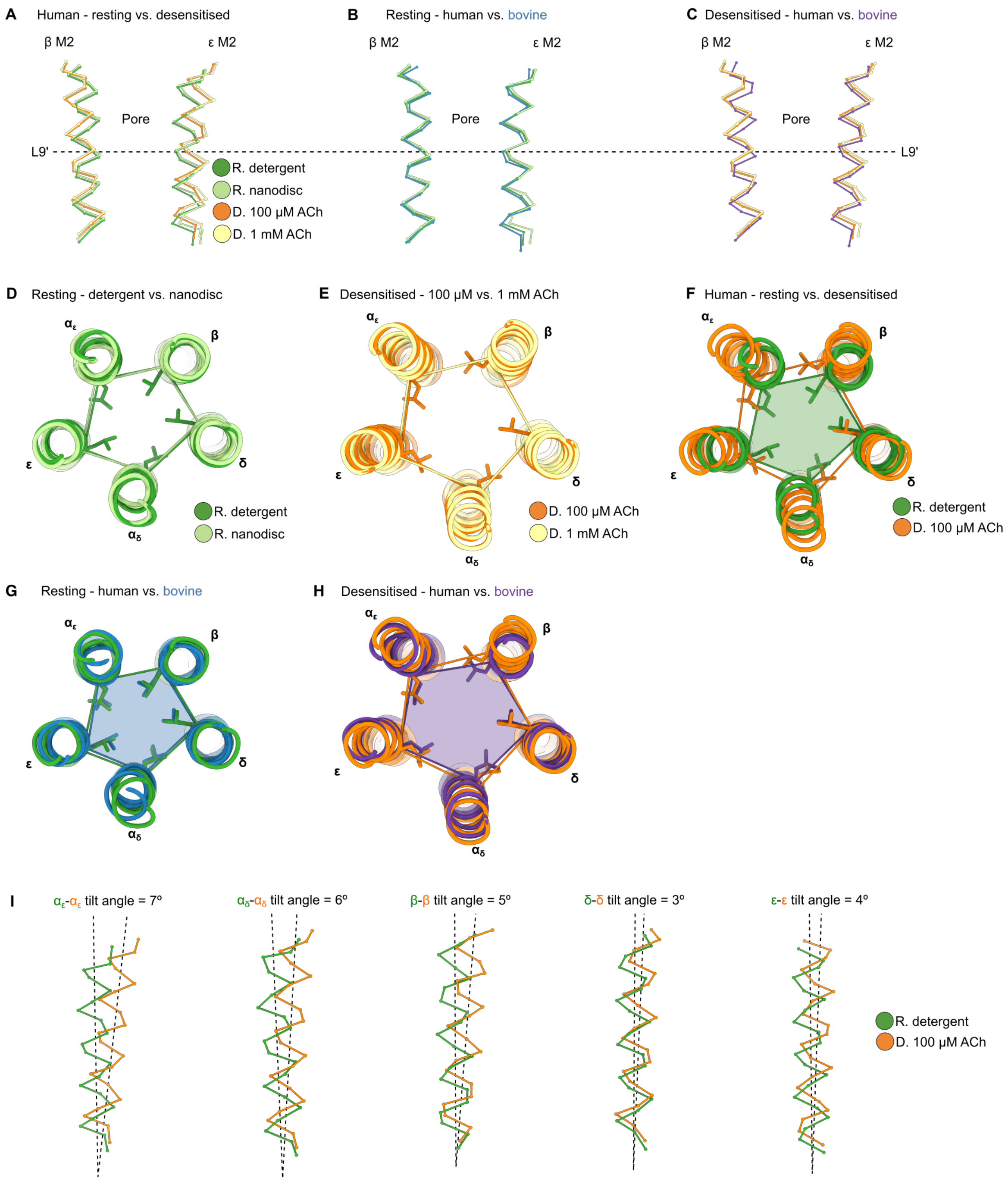


**Figure S9: Comparisons of human versus bovine AChR pore conformations.** (A) Superposition of four human receptor structures from the present study showing the cross-section at β M2 and ε M2. (B-C) Comparison of human versus bovine M2 conformations in the resting and desensitised states (PDB: 9AVV and 9AWJ)[6]. (D-E) Comparison of pore dilation at the 9’ gate for the two resting and two desensitised structures. (F) The 9’ leucine residues rotate away from the pore axis in the desensitised conformation. (G) Human and bovine resting state structures show the same level of dilation at the 9’ position, whereas (H) in the desensitised state, β subunit of the human receptor is located further away from the pore axis. (I) M2 tilt angles between resting and desensitised M2 helices were calculated with PyMOL (Version 1.20 Schrödinger, LLC.).

| **Table S3: Sequence variants identified in cases of suspected congenital myasthenic syndrome** | | |
|---|---|---|
| **Gene** | **Amino acid variant (from first codon of mature peptide)** | **RefSeq_mRNA/hgvs_DNA/hgvs_protein** |
| CHRNA1 | αI49T | NM_000079.4 (CHRNA1): c.206C>T (p.Ile69Thr) |
| CHRNA1 | αT133I | NM_000079.4 CHRNA1: c.458C>T (p.Thr153Ile) |
| CHRNA1 | αT281S | NM_000079.4 (CHRNA1): c.902C>G (p.Thr301Ser) |
| CHRNB1 | βI285S | NM_000747.3 (CHRNB1): c.923T>G (p.Ile308Ser) |
| CHRNB1 | βV296A | NM_000747.3 (CHRNB1) c.956T>C (p.Val319Ala) |
| CHRND | δD180N | NM_000751.3 (CHRND): c.601G>A (p.Asp201Asn) |
| CHRND | δL272P | NM_000751.3 (CHRND): c.878T>C (p.Leu293Pro) |
| CHRNE | εS235A | NM_00080.4 (CHRNE): c.763T>G (p.Ser255Ala) |

| **Table S4: Single channel kinetics of WT and mutant AChR expressed in HEK293T cells** | | | | | | | | | |
|---|---|---|---|---|---|---|---|---|---|
| **Mutation** | a1 | τ1 (ms) | a2 | τ2 (ms) | a3 | τ3 (ms) | a4 | τ4 (ms) | Number of patches |
| **WT adult** | 0.28 ± 0.01 | 0.14 ± 0.01 | 0.34 ± 0.02 | 1.301 ± 0.10 | 0.38 ± 0.01 | **5.58 ± 0.34** | | | 29 |
| **WT foetal** | 0.39 ± 0.04 | 0.09 ± 0.01 | 0.24 ± 0.03 | 1.71 ± 0.38 | 0.38 ± 0.02 | **11.28 ± 0.73** | | | 4 |
| **αI49T*** | 0.35 (± 0.01) | 0.17 (± 0.05) | 0.21 (± 0.01) | 1.83 (± 0.08) | 0.33 (± 0.01) | 11.77 (± 0.07) | 0.12 (± 0.02) | **56.96 (± 0.13)** | Combined from 4 patches |
| **αT133I** | 0.49 ± 0.03 | 0.40 ± 0.05 | 0.51 ± 0.03 | **1.86 ± 0.13** | | | | | 5 |
| **αT281S** | 0.39 ± 0.02 | 0.14 ± 0.01 | 0.23 ± 0.01 | 1.72 ± 0.19 | 0.38 ± 0.01 | **18.11 ± 2.43** | | | 7 |
| **βI285S** | 0.33 ± 0.03 | 0.09 ± 0.02 | 0.44 ± 0.02 | 0.44 ± 0.07 | 0.23 ± 0.04 | **1.81 ± 0.29** | | | 6 |
| **βV296A** | 0.40 ± 0.01 | 0.10 ± 0.01 | 0.33 ± 0.01 | 0.98 ± 0.10 | 0.27 ± 0.004 | **48.07 ± 2.08** | | | 5 |
| **δD180N** | 0.54 ± 0.05 | 0.24 ± 0.01 | 0.46 ± 0.05 | **1.05 ± 0.04** | | | | | 3 |
| **δL272P** | 0.41 +-/ 0.03 | 0.17 ± 0.01 | 0.23 ± 0.03 | 3.58 ± 0.62 | 0.36 ± 0.03 | **26.09 ± 3.43** | | | 5 |
| **εS235A*** | 0.42 (± 0.01) | 0.39 (± 0.05) | 0.24 (± 0.02) | 9.19 (± 0.10) | 0.34 (± 0.02) | **69.88 (± 0.07)** | | | Combined from 7 patches |
| **εD163E + εD173F** | 0.39 ± 0.02 | 0.11 ± 0.01 | 0.32 ± 0.04 | 1.07 ± 0.21 | 0.28 ± 0.05 | **4.17 ± 1.05** | | | 6 |

*Due to low activity in individual patches, defined bursts from multiple patches were combined and fitted to multiple exponential functions. SEM for these entries represent fitting error.

**Table S5: CMS variants affecting AChR kinetics**

| SCCMS | FCCMS |
|---|---|
| **αI49T** | **αT133I** |
| αG153S/A[8] | αV132L |
| αV156M | αV188M |
| αN217K | αF256L |
| αV249F/G[9] | αV285I |
| αT254I | αG378D[10] |
| αS269I | βP248L[11] |
| αG275V[12] | **βI285S** |
| **αT281S** | δE59K |
| αC418W | δD140N[13] |
| βV229F | **δD180N** |
| **δL272P** | δP250Q |
| βL262M | εT38K |
| βT265S | εW55R |
| βV266M/A[14] | εP121L |
| **βV296A** | εD175N |
| δI261T[15] | εN182Y |
| δS268F | εE184K |
| δL273F[16] | εR218W[17] |
| εL221F [18] | εA411P |
| **εS235A** | εc.1254ins18 |
| εP245L[19] | εN436del |
| εV259L/F[20] | |
| εT264P | |
| εV265A | |
| εL269F*[21] | |
| εS278del[22] | |

*Table was updated from a 2015 review article[23]. Additional CMS variants are included with reference, variants from this study are coloured.

**A separate εL269F variant arose from an unrelated family.

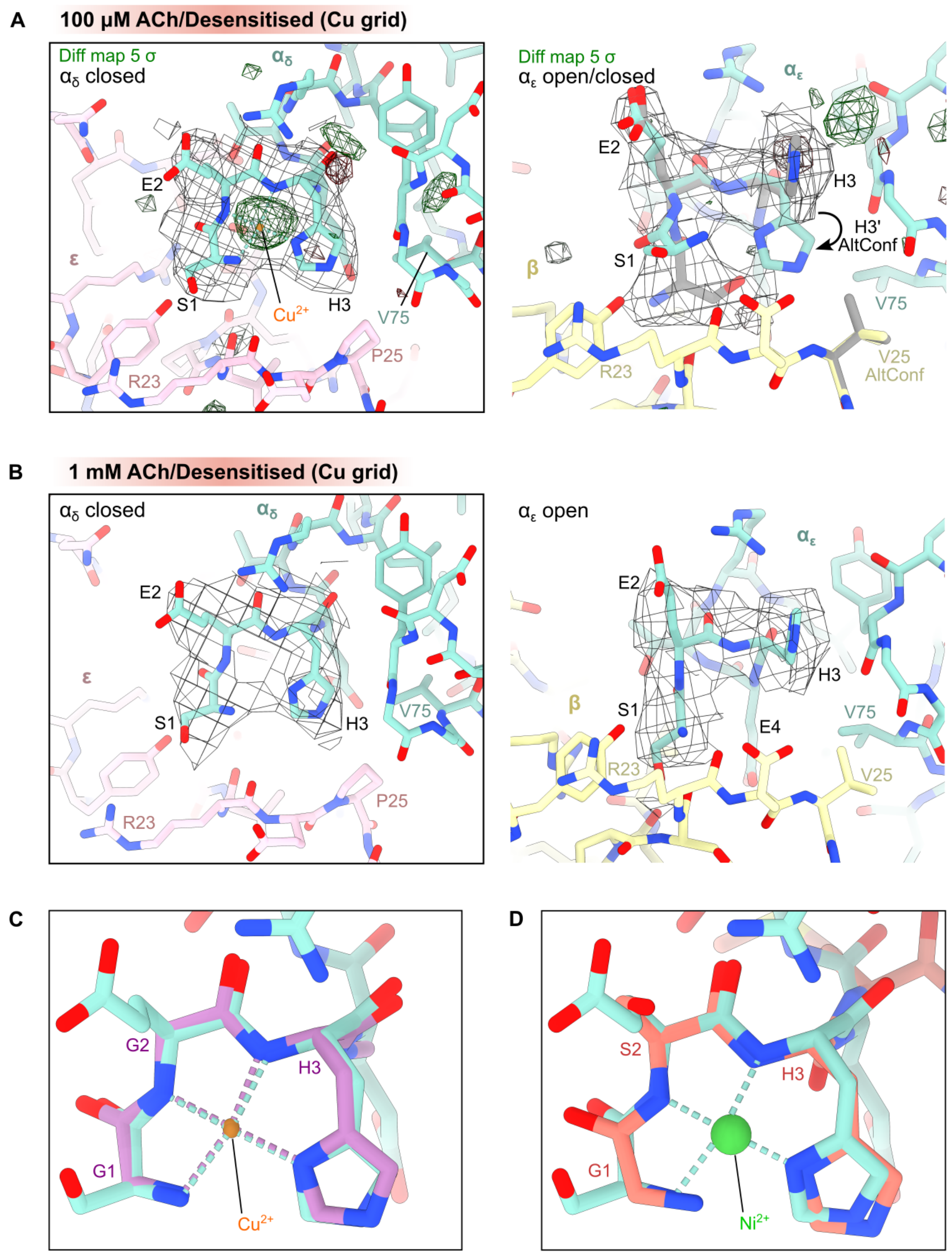


**Figure S10: Comparison of amino terminal $Cu^{2+}$ and $Ni^{2+}$ (ATCUN) binding motifs.** (A) The desensitised AChR structure in 100 µM ACh contains closed ATCUN motif (X1-X2-H3) at $\alpha_\delta$-ε site and open ATCUN at $\alpha_\varepsilon$-β site. Alternative conformations were observed for $\alpha_\varepsilon$H3 and βV25. Coulomb potential maps are shown in grey, mask-normalised $F_o$-$F_c$ difference maps highlighting non-protein features are shown in green (positive density) and red (negative density), calculated with Servalcat (see Methods). (B) The second desensitised structure (in 1 mM ACh) adopts closed and open conformations at corresponding sites. (C) Overlay of copper-bound glycylglycyl-L-histidine-N-methyl amide structure[24] and $\alpha_\delta$ ATCUN motif of resting state nanodisc AChR structure, RMSD=0.22 Å. Structure of peptide was obtained from the Cambridge Structural database[25]. (D) Overlay of ATCUN motifs on nickel-bound c-Src-SH3 domain (PDB:4RTZ) and resting state AChR, RMSD=0.31 Å[24,26].

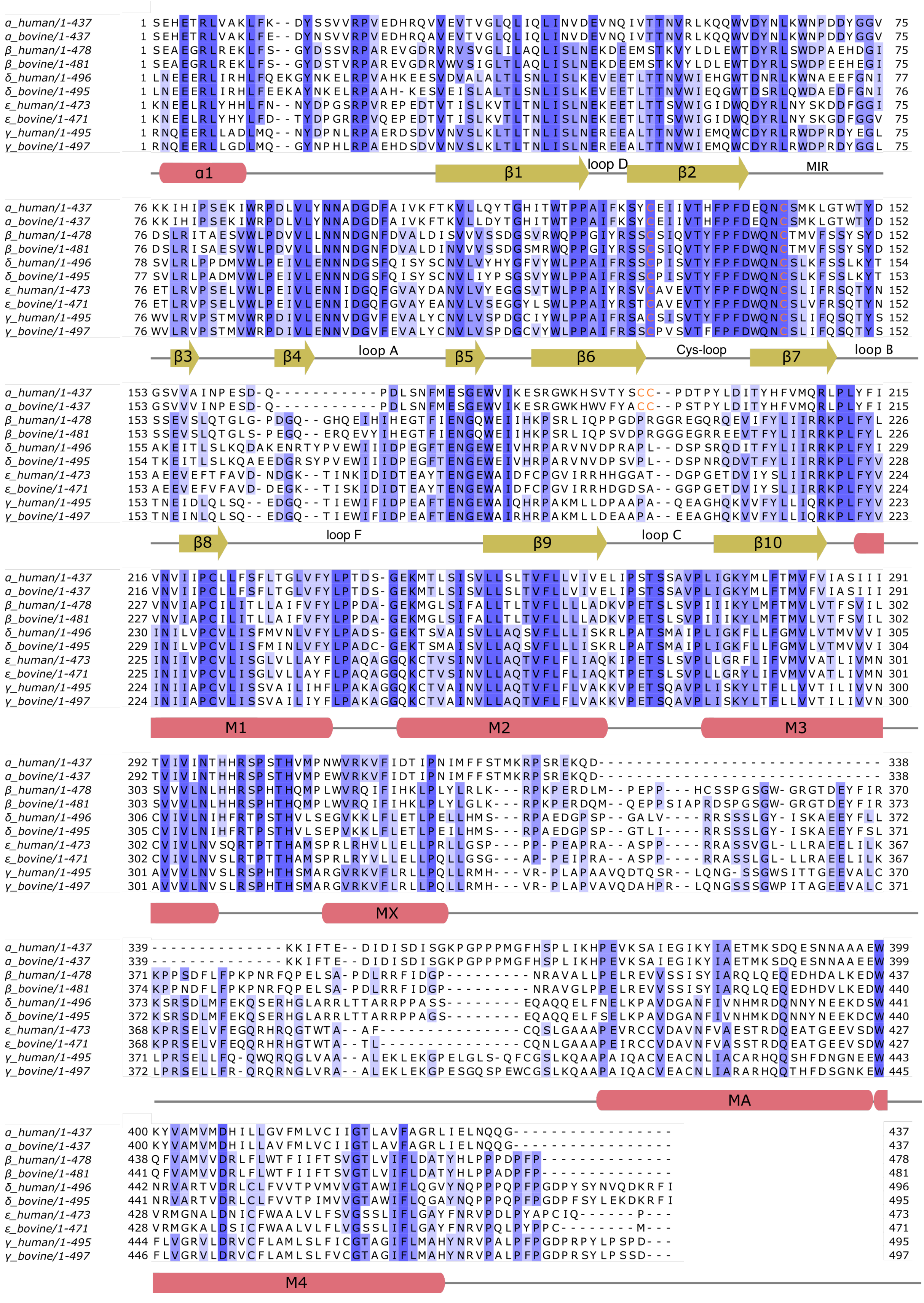


**Figure S11**: Sequence alignment of human versus bovine muscle-type AChR subunits. Pipes and arrows indicate approximate positions of α-helices and β-strands[7]. UniProt sequences (P02708, P02709, P11230, P04758, Q07001, P04759, Q04844, P02715, P07510, P13536)

were aligned with Tcoffee in Jalview[27]. Signal peptides were removed from residue numbering according to positions in original publications[28-33]. Cysteines involved in disulphide bonding are coloured orange.
